# Retinoic acid, an essential component of the RP organizer, promotes the spatio-temporal segregation of dorsal neural fates

**DOI:** 10.1101/2024.03.17.585392

**Authors:** Dina Rekler, Shai Ofek, Sarah Kagan, Gilgi Friedlander, Chaya Kalcheim

**Author notes:** To whom correspondence should be addressed. Emails: Dina Rekler,; Gilgi Friedlander, Shai Ofek, Sarah Kagan, Chaya Kalcheim.

## Abstract

Dorsal neural tube-derived retinoic acid promotes the end of neural crest production and transition into a definitive roof plate. Here we analyze how this impacts the segregation of central and peripheral lineages, a process essential for tissue patterning and function. Localized in-ovo inhibition of retinoic acid activity followed by single cell transcriptomics unraveled a comprehensive list of differentially expressed genes relevant to these processes. Importantly, progenitors co-expressed neural crest, roof plate and dI1 interneuron markers indicating a failure in proper lineage segregation. Furthermore, we found that separation between roof plate and dI1 interneurons is mediated by Notch activity downstream of retinoic acid, highlighting their critical role in establishing the roof plate-dI1 boundary.

Within the peripheral branch, where absence of retinoic acid resulted in neural crest production and emigration extending into the roof plate stage, sensory progenitors failed to separate from melanocytes leading to formation of a common glia-melanocyte cell with aberrant migratory patterns. Together, we uncover and characterize a molecular mechanism responsible for segregation of dorsal neural fates during development.

## Introduction

The separation between central and peripheral branches of the nervous system (CNS and PNS), takes place in the dorsal domain of the neural tube (NT), offering a prototypic example of lineage segregation during development. This domain sequentially generates neural crest cells (NC), progenitors of the PNS, followed by the definitive roof plate (RP) of the CNS, which is bordered ventrally by dorsal interneurons ^1–3^.

NC progenitors undergo a Bone Morphogenetic Protein (BMP)-dependent epithelial to mesenchymal transition (EMT), exit the NT and generate neurons and glia of peripheral ganglia, as well as melanocytes^4–6^. RP progenitors originate ventral to the premigratory NC, relocate ventro-dorsally as a result of continuous NC emigration, and reach the midline region of the NT upon completion of NC departure^7,8^. In contrast to NC, RP cells change their state of competence and become refractory to BMP signaling despite continuous ligand synthesis^9^. They progressively exit the cell cycle and act as a dorsal organizer, providing BMP and Wnt ligands required for interneuron development^3,9–14^.

Hence, our understanding of cellular states/fates and the mechanisms of transitions progressively evolves. However, the mechanisms responsible for lineage separation of NC-derived cell types, of NC from RP, and RP from adjacent dorsal interneurons, remain challenging questions at the experimental level, given the rapid temporal dynamics of these processes.

A prerequisite for tackling the transition between NC and RP is to know how NC production ceases. We found that dorsal NT-derived retinoic acid (RA) ends the period of NC production and emigration via inhibition of BMP/ Wnt activities. This is accounted for by restricted and dynamic expression of BMP antagonists in the dorsal NT and by a network of specific downstream effectors. In absence of RA activity, NC emigration extended into the RP period, and dI1 interneurons also invaded the RP territory. Hence, the temporal and spatial segregation of dorsal lineages, as well as the structural integrity of the RP were abnormal^15^. Furthermore, we showed that both gain and loss of Notch function altered the balance between RP and dI1 interneurons, suggesting this pathway is required for setting the RP-dI1 boundary^16^. A refined analysis of the above processes and their integration into a comprehensive molecular network are still lacking.

By inhibiting RA activity locally followed by single-cell RNA-seq we here uncover: first, a large set of differentially expressed genes compared to normal embryos. Second, lack of RA prevented the separation of NC, RP, and dI1 interneuron fates, leading to combinatorial co-expression of lineage markers within single cells. Third, Notch activity downstream of RA mediates the separation between RP and dI1 interneurons.

Likewise, in the absence of RA, the extended production and emigration of NC cells led to PNS progeny in which separation of sensory glia and melanocytes, thought to derive from a common progenitor ^17–20^, was hindered. As a result, single cells with glia-melanocyte traits were apparent, that exhibited atypical migratory patterns. These findings show that RA is responsible for the segregation of numerous dorsal neural fates in a cell type-specific manner, highlighting the significance of RA and related signaling pathways in normal fate acquisition and tissue patterning.

## Results

### Single-cell transcriptomics identifies a developmental dynamics of cell types in the dorsal NT

To enrich for dorsal NT cell types, quail E2.5 NTs were electroporated at the flank region of the axis with Msx1-Citrine^21^ combined with either a control vector, or RARα403, a dominant negative RA receptor. For each group, ten NTs with associated mesoderm were mechanically microdissected at E4 and single cell suspensions prepared. Fluorescent cells were then sorted and subjected to droplet-based scRNA-seq (10x Genomics Chromium) to generate a gene expression profile of control and treated dorsal NT cells (Fig. 1A-D). In total, 17,549 cells were sequenced. After applying quality filters (see Methods), a dataset of 14,566 cells was retained for further analysis (5,890 control and 8,676 treated cells, which comprise 86.6% and 80.7% of the initial number of cells, respectively).

**Figure 1.**
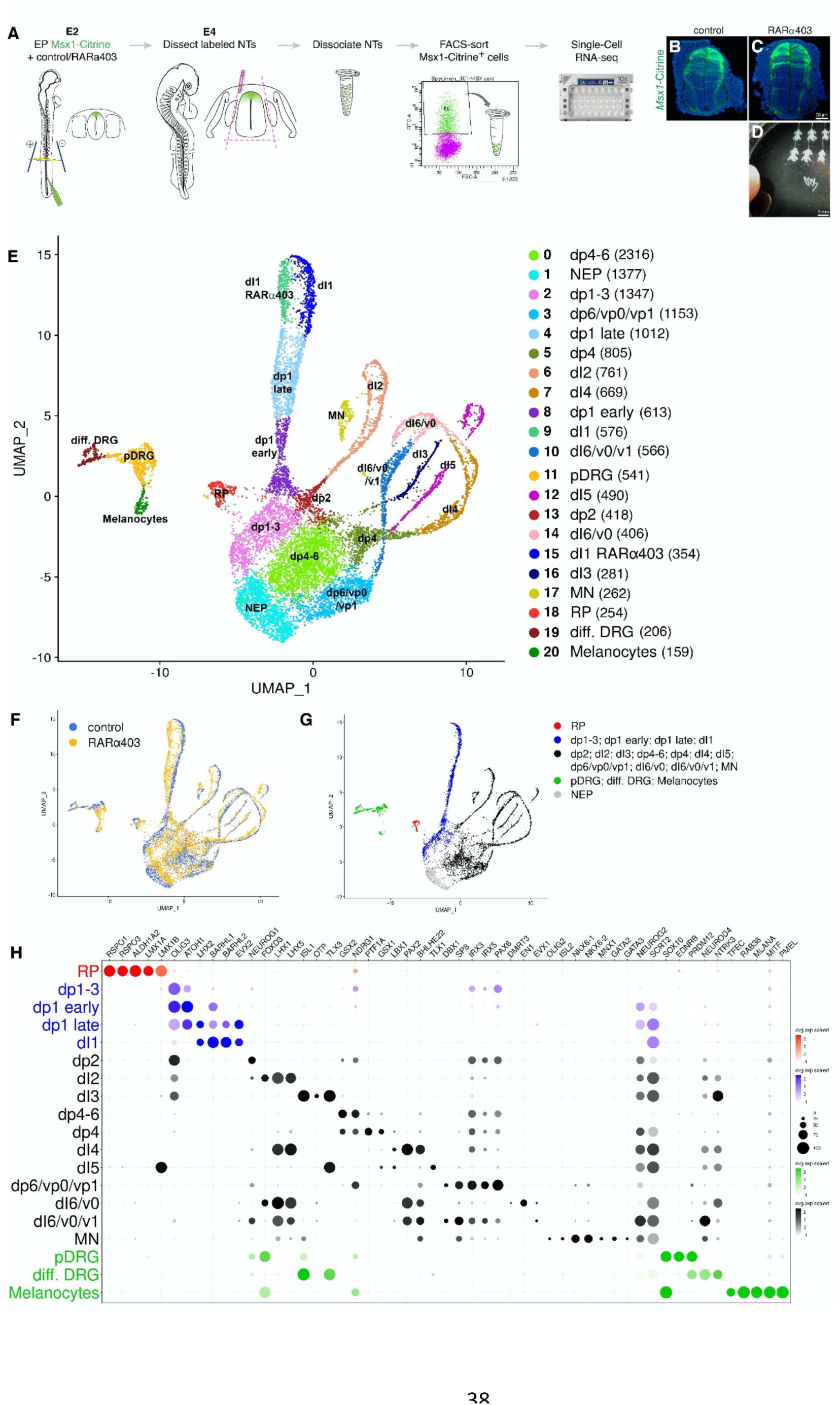
Assignment of transcriptomes to cell identities. A) Schematic representation of the experimental design for isolating single dorsal NT cells. B,C) NTs were electroporated at E2.5 and microdissected at E4, showing a similar pattern of Msx1-Citrine labeling between control and RARα403-treated embryos. D) An example of embryos harvested for processing (top 2 rows) and of isolated NTs (bottom). E) UMAP plots of control and treated samples, colored according to the cluster. The number of cells in each cluster is indicated in parentheses. F) UMAP of the entire dataset colored by sample. G-H) Cluster identification based on expression of selected marker genes in the control sample. In the dot plot (H), the average expression level and percentage of expressing cells per gene are illustrated by the color intensity and the diameter of the dot, respectively. Abbreviations, dI, dorsal interneurons; diff. DRG, differentiated dorsal root ganglion neurons; dp, dorsal interneuron progenitors; EP, electroporation; MN, motoneurons; NEP, neuroepithelial progenitors; NT, neural tube; pDRG, dorsal root ganglion progenitors; RP, roof plate. Scale bar, 50 μm

#### Assignment of transcriptomes to cell identities

Unsupervised clustering, identified 21 distinct cell clusters, which were visualized using Uniform Manifold Approximation and Projection (UMAP) (Fig. 1E). Most of the clusters contained similar numbers of control and treated cells (Fig. 1F and Supplem. Fig1A). The clusters were allocated to different cell types based on combinatorial expression of a selected list of established genes in the control sample (Fig. 1G-H) ^22,23^.Mainly dorsal NT cell types (RP, dI1-6 interneurons) and NC-derived DRG and melanocytes^24^ were identified but not NC derivatives bearing a more ventral character.

The latter is consistent with the timing of electroporation that attained only the later emigrating NC progenitors^8^. Clusters 3 and 14 comprised, respectively, progenitors or neurons expressing mixed dI6/v0/v1 characteristics perhaps due to poorly resolved transcriptomes, and cluster 17 contained motoneurons. Although significant enrichment of dorsal NT cells was apparent, the presence of various ventral cell types suggests some leaky expression of Msx1-Citrine at early transfection.

#### A map of time and space

The resulting UMAP bears the shape of an “octopus-like” structure with a head and emerging arms. Cell cycle analysis showed that the octopus head (clusters 0,1,2,3) is mainly composed of immature proliferating neuroepithelial progenitors encountered either in G2/M or the S-phases of the cell cycle (Sup. Fig. 1A). Consistently, cell cycle-associated genes such as *BUB1* and *CDK1* (Supplem. Fig. 1C), as well as the neural progenitor marker *SOX2*, etc. were expressed. Whereas cluster 1 was composed of neuroepithelial progenitors (NEP) lacking a specific identity, clusters 0, 2 and 3 contained defined dorsal interneuron precursors. Clusters 8, 13 and 5, connecting the head of the octopus with its arms exhibited, among others, a specific expression of early neuronal progenitor genes characterized by the Notch ligands *DLL1* and *JAG2* (Suppl. Fig. 1C). Furthermore, the neuronal specification markers *NEUROD1* and *NEUROG2* decorated the proximal portions of all arms, including young E4 DRG neurons (cluster 19) (Suppl. Fig. 1C). Finally, the distal portions of the arms expressed neuronal markers such as *ELAVL2*, transcription factor *EBF1*, axonal glycoprotein contactin 2 (*CNTN2*), axonal guidance receptor *ROBO3* (Suppl. Fig. 1C), as well as *NEFM, NTRK3, NHLH2, SEPTIN3, ONECUT1* and *2,* etc. A majority of cells in the latter category were in the G1 phase of the cell cycle (Suppl. Fig. 1B,C). Thus, the head-to-arm dimension reflects a temporal sequence of neural progenitor differentiation, and highlights the co-existence of immature precursors and differentiated neurons in the E4 NT.

Likewise, a spatial map of cell types was deduced along the left-to right direction of the UMAP (Sup. Fig. 1C,D), with dorsal cell types expressing *FOXD3* in DRG, *RSPO1* in RP, *WNT1* in RP and dp1-3 progenitors, *OLIG3* in dp1-3 cells, dI2 and dI3, *GSX2* in dp4-6, graded *LGR5* in dp6 to dp4-6, *LHX2* in dI1, *FOXD3* in dI2, *TLX3* in dI3/5 and *PAX2* in dI4/6 (Suppl. Fig. 1E). Hence, the left-to-right dimension of the UMAP corresponds to a dorso-ventral distribution of cell types in the E4 NT.

### RA signaling prompts fate segregation of dorsal NT cell types

#### Analysis of the normal RP predicts novel gene expression patterns

Next, we focused on the development of the RP. Sup. Fig. 2 illustrates the profile of the most significant genes expressed in control RP compared to remaining dorsal clusters. Whereas the presence of previously studied RP-specific genes, e.g; *LMX1A, LMX1B* ^25,26^*, GDF7* ^27^*, BMP4, BMP5, BMP7, BAMBI, ALDH1A2, WNT1* and *WNT3A* ^7,15,16^ was confirmed, novel unique genes were unraveled. These include *LMO7*, olfactomedin3 (*OLFML3*), calponin 2 (*CNN2*), chondromodulin (*CNMD*), the tumor supressors *CD99* and *CD82*, the metallopeptidases *ADAMTS8* and *ADAMTS3, WNT9A, UNC5C, TM6SF1, SPON1, RSPO3, KCNIP4,* etc. An additional array of mRNAs, relatively enriched in RP vis-à-vis dorsal interneurons and/or their progenitors (e.g; *LOXL1, LOXL3, LAMC1, CTBP2, DRAXIN, ZIC1-3*, etc), was also evidenced. Together, these new RP genes provide a valuable resource for further functional analysis.

#### Differential gene expression in RP of control and RARα403-treated NTs

Next, we addressed the effects of inhibiting RA signaling on RP development. Changes in gene expression between control and RARα403-treated samples revealed profound effects in four main categories: 1) Signaling pathways, 2) Transcription factors and binding proteins, 3) Extra-cellular matrix (ECM), EMT and migration, and 4) Axonal growth and guidance. Validation of representative genes by in-situ hybridization (ISH) and reporter assays confirmed the bioinformatic data (Figs 2,3, asterisks).

**Figure 2.**
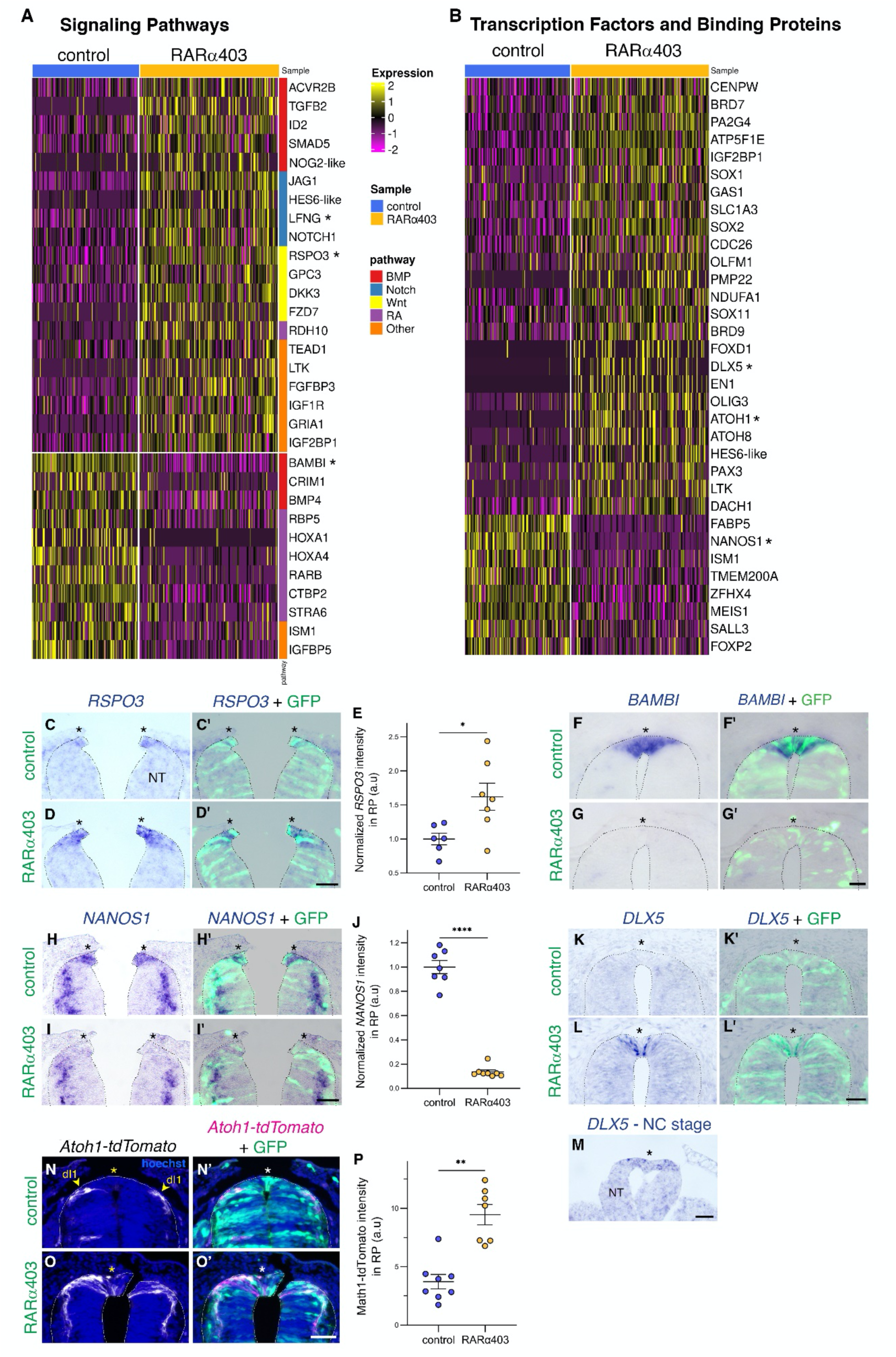
Differential gene expression in RP reveals changes in signaling pathways and transcription factors. A,B) Heatmaps of differentially expressed genes in response to RARα403 treatment in the RP cluster. Shown are selected genes with a minimum linear fold change of ±1.3 and adjusted p-value <0.05. Signaling pathway genes are depicted in (A) and transcription factors and binding protein categories in (B). Genes validated in vivo are marked with an asterisk. C-M) ISH on embryos electroporated with with GFP along with control PCAGG or RARα403 at E2.5 and analyzed at E4. The RP is marked by asterisks. C-E) ISH for *RSPO3*, showing upregulation in treated embryos, N = 6,7 embryos for control and treated groups, respectively. E) data quantification. F-G) ISH for *BAMBI*, showing downregulation in treated embryos (see Rekler et al, 2023 for details). H-J) ISH for *NANOS1,* showing downregulation in the RP of treated embryos, N = 7,8 embryos for control and treated groups, respectively. J) data quantification. K-M) *DLX5 mRNA* is expressed in premigratory NC (M), downregulated in control RP (K,K’), yet shows extended expression in treated RP (L,L’). N = 6,6,3 for control, RARα403 and NC groups, respectively. N-P) Electroporation at E2.5 with GFP and Atoh1-tdTomato, along with control PCAGG or RARα403, followed by fixation at E4. The Atoh1-tdTomato reporter labels dI1 cell bodies and processes. Yellow arrowheads mark the location of dI1 cells ventral to RP in control embryos. Note, however, invasion of the treated RP by cells with reporter activity. P) data quantification N = 8,7 embryos for control and RARα403 groups, respectively. Abbreviations, a.u, arbitrary units; dI1, dorsal interneurons1; NC, neural crest; NT, neural tube; RP, roof plate. *p < 0.05, **p < 0.005, ****p < 0.0001, Welch’s t-test (E,J) and Mann-Whitney test (P). Scale bar, 50 μm.

**Figure 3.**
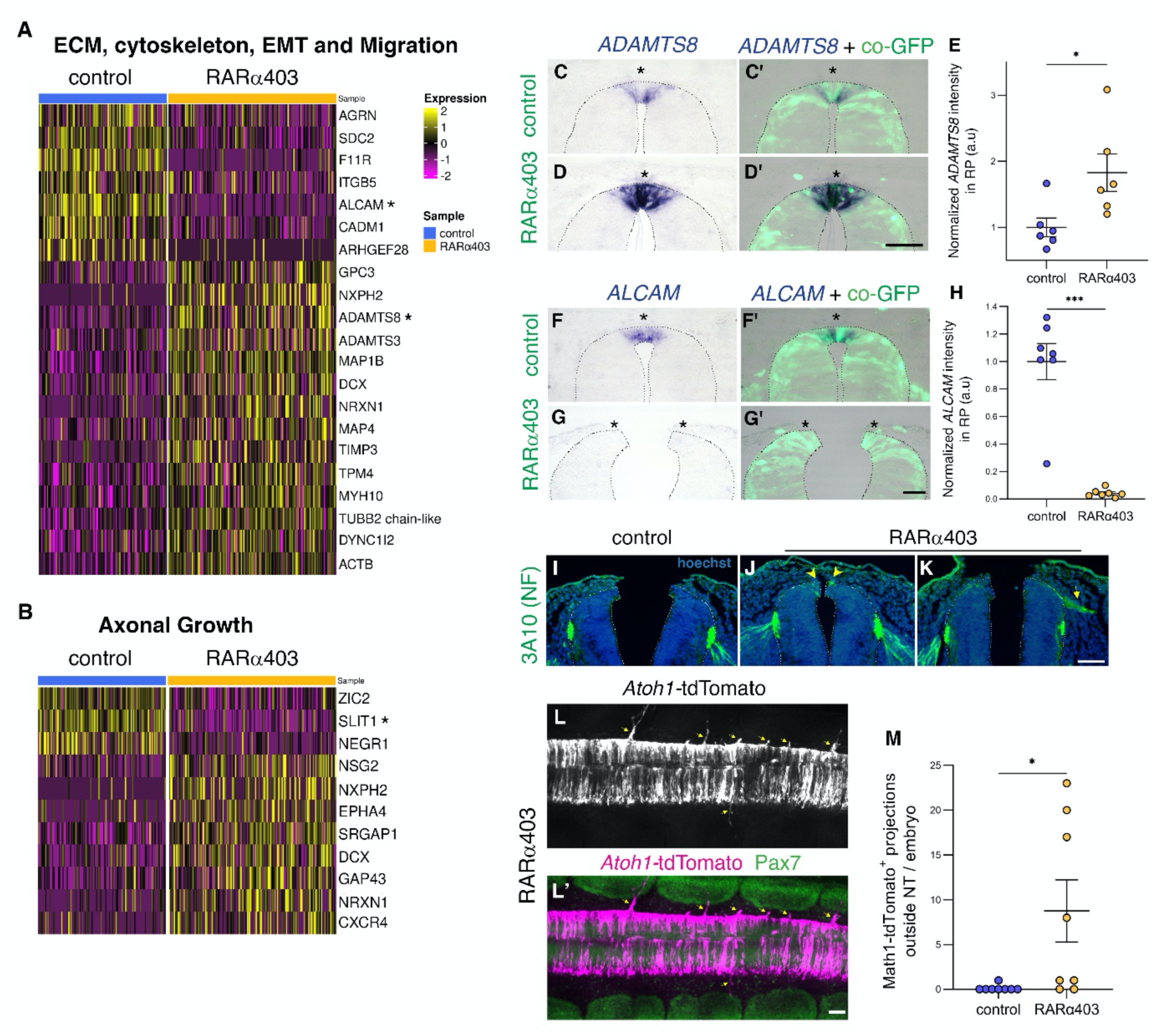
Differential gene expression in RP reveals changes in adhesive properties and axonal growth. A,B) Heatmap of differentially expressed genes in response to RARα403 treatment in the RP cluster. Shown are selected genes, with minimum linear fold change of ±1.3 and adjusted p-value <0.05 involved in ECM, cytoskeleton, EMT and migration (A) and axonal growth and guidance (B) categories are shown. Genes validated in vivo are marked with an asterisk. C-M) In-vivo validation on embryos electroporated with GFP along with control PCAGG or RARα403 at E2.5 and analyzed at E4. The RP is marked by asterisks. C-E) ISH for *ADAMTS8*, showing upregulation in treated embryos, N = 6,6 embryos for control and treated groups. E) data quantification. F-H) ISH for *ALCAM*, showing downregulation in treated embryos, N = 7,7 embryos for control and treated groups. H) data quantification I-K) Staining for neurofilament protein (3A10 antibody), showing the presence of immunoreactive cells in treated RPs (arrowheads in J) and of projections extending from the dorsal NT outward (arrow in K); N = 9,9 embryos for control and RARα403 groups. L-M) Embryos were electroporated as above, along with the Atoh1-tdTomato reporter. Anti-Pax7 was used to stain dorsal NT and somites. Note that Atoh1-tdTomato^+^ axonal projections abnormally extend from the dorsal NT into the mesoderm (arrows). M) data quantification. N = 8,8 embryos for controls and RARα403. *p < 0.05, ***p < 0.001, Mann-Whitney test. Scale bar, 50 μm.

As previously reported, BMP and Wnt activities are maintained in RA-depleted dorsal NTs as opposed to their normal downregulation in RP when compared to premigratory NC ^15^. Consistently, numerous Wnt pathway genes were upregulated, including *RSPO3*, and so were the positive BMP effectors *ID2* and *SMAD5* (Fig. 2A)^28,29^. Simultaneously, BMP inhibitors like *BAMBI* and *CRIM1* ^30,31^, as well as *BMP4* itself were reduced. Together, a balance between positive and negative BMP pathway genes is likely to account for the measured upregulation in factor activity ^9,15^.

Alterations in RA pathway genes were anticipated upon repression of RA signaling. Indeed, this negatively impacted various pathway genes, including the RA receptor *RARB*, the retinol binding protein *RBP5*, *STRA6*^32^ and *CTBP2*^33^, though we observed an upregulation of *RDH10*, a RA biosynthetic enzyme ^32^. Notably, all genes related to the Notch pathway were upregulated, hinting at a possible repression of RA on Notch signaling (Fig. 2A and see also Fig. 5).

Many transcription factors were affected in RARα403-treated NTs (Fig. 2B). These included upregulated *SOX2*, consistent with RA acting as a differentiation factor in absence of which cells preserve a progenitor-like state. Notably, the RNA-binding protein *NANOS1* was downregulated (Fig. 2B, H-J). NANOS1 was reported to be involved in germ cell development, hippocampal development and neurogenesis ^34,35^. In our RNA-seq, it is a novel RP marker of unknown function.

Most importantly, a distinct set of upregulated transcription factors consisted of NC (e.g. *DLX5*) and dI1-specific (*OLIG3, ATOH1, ATOH8*) genes whose expression in treated vs. control RP was validated in vivo by ISH and by a reporter assay for *ATOH1* (Fig. 2K-M, N-P). This confirms and further extends our findings that RP, NC and dI1 interneurons fail to segregate in absence of RA signaling ^15^.

Inhibition of RA activity leads to an extended period of cell emigration from the dorsal NT subsequent to completion of the NC stage^15^. Therefore, we expected a significant impact on genes involved in cytoskeletal function, EMT and migration. Indeed, metalloproteases such as *ADAMTS8, ADAMTS15* and *TIMP3*, microtubules and myosin-associated genes such as *MAP4, TUBB2, TUBB3, MYH9* and *10*, etc. as well as genes involved in neuronal precursor migration such as *DCX* were upregulated (Figs 3A,C-E and Sup. Fig 3C). Reciprocally, genes involved in cell adhesion such as *ALCAM* were downregulated (Fig. 3A,F-H).

Lastly, a set of RP genes affected by RARα403 is related to axonal growth and guidance (Fig. 3B, Sup. Fig 3B). For instance, the axonal repellent *SLIT1*(Rekler and Kalchein, 2022), the chemokine receptor *CXCR4*, also involved in cell migration^36^, the *EPHA4* and *EPHA3* receptors ^37^, *GAP43* which is involved in axonal growth and guidance, etc. Consistently, we find that RA is important for the correct patterning of neuronal cell processes, as neurofilament-expressing fibers were abnormally detected in the treated RP. In addition, atypical neuronal projections extended out of the dorsal NT (Fig. 3I-K) and were shown by a Atoh1 reporter, to correspond to dI1 interneurons (Fig. 3L-M), that normally project dorso-ventrally within the NT ^38^. These results extend our previous findings showing an infiltration of BarHL1-positive dI1 interneurons into the RP in the absence of RA signaling ^15^. The specific molecular mechanism accountable for the observed phenotypes among the aforementioned altered pathways is yet to be explored.

#### Single RP cells co-express RP, NC and dI1 traits

We previously documented the significance of RP-derived RA signaling for the end of NC production, an effect that included the downregulation of NC-specific genes such as *FoxD3*, *Snai2* and *Sox9* in the dorsal NT and consequent lack of NC EMT ^15^. Furthermore, extended misexpression of these genes into the RP stage prevented the timely upregulation of *ALDH1A2* mRNA. Using a RP-specific *ALDH1A2* enhancer, we now confirm and extend these results to show that upon late transfection of NC-specific genes, enhancer activity is reduced (Sup. Fig. 4). Thus, NC and RP genes stand in a mutually repressive interaction along a temporal axis, enabling a transition between the above states.

As shown in Fig. 2, lack of RA activity interfered with the above transition, as genes specific for RP, NC and also dI1 genes were all evident in the RP cluster (see also^15^). It was therefore important to understand the effect of RA inhibition on the RP at single cell resolution. To this end, control and treated RP cells were re-clustered into four sub-clusters, based on similarity in gene expression (Sup. Fig. 5). Clusters 0 and 3 were primarily composed of control cells, showing few differences in gene expression. On the other hand, clusters 1 and 2 consisted mainly of RARα403-treated cells, with cluster 2 demonstrating an intermediate expression profile partially similar to that of the control cells, perhaps due to a lower transfection efficiency. In contrast, cluster 1, while retaining expression of the RP markers *RSPO1* and *RSPO3*, differed significantly from the control clusters, encompassing the majority of the effects unveiled in the differential expression analysis (Fig 2,3, Sup. Fig. 3). Particularly, genes associated with both premigratory NC (*SNAI2*, *DLX5*, *SOX9*, *NOG*) and dI1 (*ATOH1*, *OLIG3*, *ATOH8*, *SOX9*) fates were highly expressed within this cluster, hinting at a possible failure of fate segregation at the single-cell level.

Next, we examined the possibility of co-expression of RP, NC and dI1 genes in individual RP cells of treated embryos. To this end, cells with a minimum of two reads per gene were considered positive, and the co-expression of different gene pairs was analyzed. Figure 4 demonstrates that while control RP clusters contained few or no double-positive cells, a significant proportion of the treated RP cells co-expressed the RP marker *RSPO1* together with either NC (e.g. *SNAI2*) or dI1 (e.g. *ATOH1* or *OLIG3*) markers. Remarkably, co-expression of NC together with dI1 markers was detected as well (e.g. *SNAI2*-*ATOH1* and *DLX5*-*OLIG3*). Co-expression of *RSPO1* and *RSPO3* served as positive control. As expected, these genes were co-expressed both in the control and in the treated RP cells. These findings, in conjunction with additional gene pairs tested, are summarized in Fig. 4B and were further validated in-vivo using combinations of ISH for *RSPO1*, immunostaining for SNAI2 and BarHL1, and a reporter assay for *ATOH1* (Fig 4C-H”). Collectively, these results suggest that at some time during development, individual dorsal NT cells have at least three potential fates – NC, RP and dI1. Here we show that normal segregation between these fates, both in time and space, is regulated by RA.

**Figure 4.**
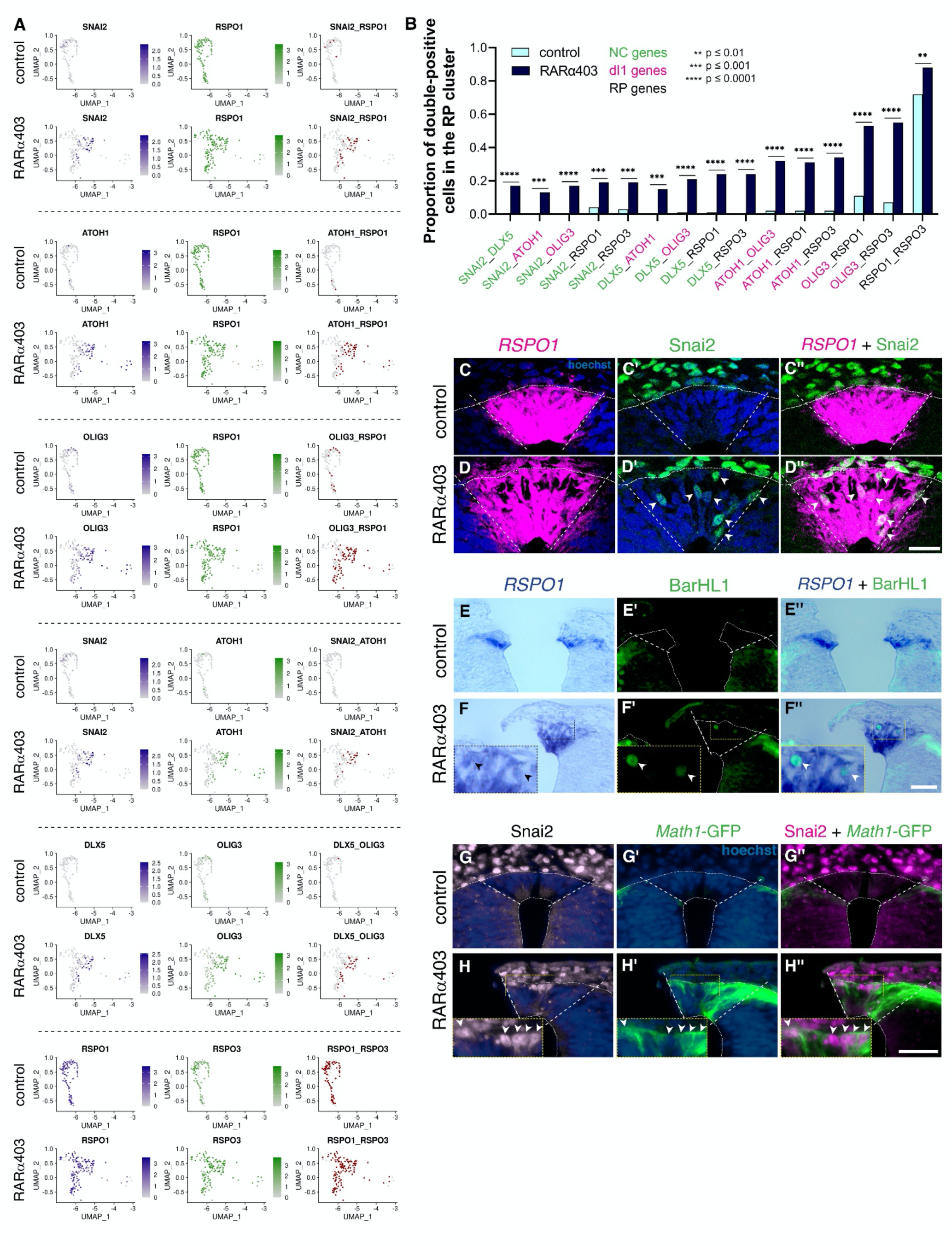
Single RP cells co-express RP, NC and dI1 traits. A) Co-expression of NC, dI1 and RP genes in the RP cluster visualized on UMAPs of control and RARα403 samples. Cells which had at least two reads/cell of the genes of interest are considered double-positive and stained red in the right column. Note the negligible proportion of double-positive cells in all control samples, as opposed to a significant amount in the treated ones. The co-expression of *RSPO1* and *RSPO3* (bottom) served as a positive control. B) Testing for the co-expression of an array of gene pairs characterizing RP, NC and dI1 interneuronal identities (see Methods). **p < 0.01, ***p < 0.001, ****p < 0.0001. C-H) In vivo validations of gene co-expression on embryos electroporated with control PCAGG or RARα403 at E2.5 and analyzed at E4. C-D) Immunostaining for SNAI2 combined with fluorescent ISH for *RSPO1*. Dashed lines mark the RP domain. Note the presence of SNAI2^+^*RSPO1*^+^ cells in the RP (arrowheads in D’,D”). N = 8,8 for controls and RARα403. E-F) Immunostaining for BarHL1 combined with ISH for *RSPO1*. Note the presence of BarHL1^+^*RSPO1*^+^ cells in the RP (arrowheads in F”). N = 6,6 for controls and RARα403. G-H) Embryos were electroporated as above along with a Atoh1-GFP reporter for dI1 labeling and immunostained for SNAI2. Note double-positive cells in the RP (arrowheads in H’’). N = 6,9 for controls and RARα403, respectively. Abbreviations, dI1, dorsal interneurons 1, NC, neural crest, RP, roof plate. Scale bar, 50 μm.

**Figure 5.**
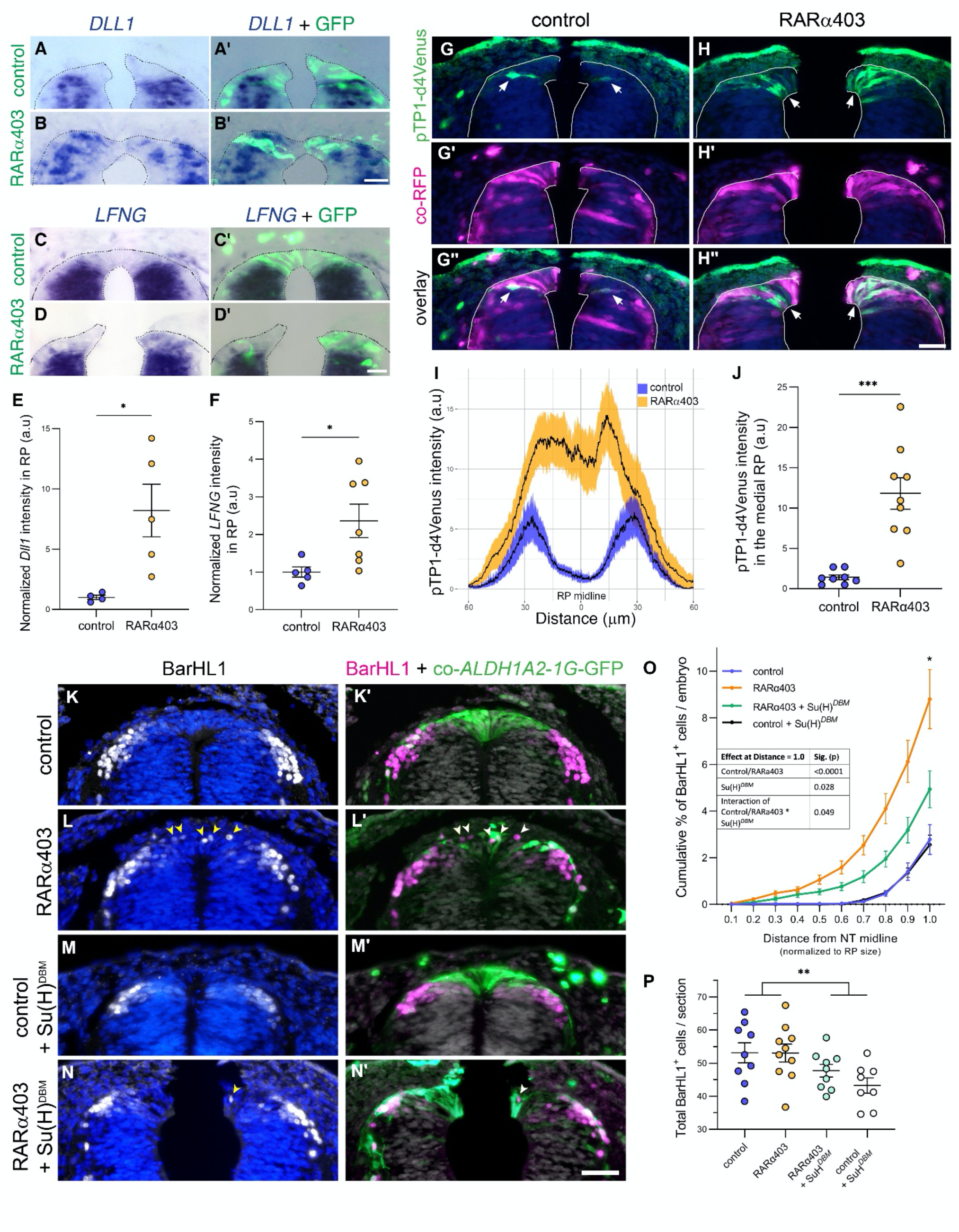
Segregation of RP from dI1 interneurons requires inhibition of Notch signaling by RA. Embryos were electroporated with control PCAGG or RARα403 at E2.5 and analyzed at E4. A-F) ISH of *DLL1* (A,B,E) and *LFNG* (C,D,F), showing positive gene expression ventral to the RP in controls, and expansion into the RP domain in treated embryos. N = 4,5 (*DLL1*) and 5,7 (*LFNG*) embryos for control and RARα403 groups, respectively. G-J) Embryos were electroporated as above, along with RFP and the pTP1-d4Venus reporter to monitor Notch activity. Arrows in G-H” mark pTP1 signal, showing a notable expansion into the RP domain of treated embryos and no signal in the RP of controls. I) Quantification of signal distribution in RP and adjacent interneurons. J) Quantification of pTP1 signal intensity in the medial 30 μm of the dorsal NT. K-P) Embryos were electroporated as above, with the addition of either a control vector or Su(H)*^DBM^*under the control of a RP-specific *aldh1a2* 1G enhancer. GFP driven by the same enhancer served as an electroporation control. Arrowheads in L,L’,N,N’ mark BarHL1^+^ cells inside the RP domain. Note the reduction of BarHL1+ cells in RP between RARα403 alone (L) and RARα403 combined with Su(H)*^DBM^* (N). O) Quantification of the percent distribution of individual BarHL1^+^ cells located inside the RP as a function of distance from the dorsal midline, normalized to RP size. Two-way ANOVA was performed on the data at Distance = 1.0, showing a significant effect of Su(H)*^DBM^* in reducing the phenotype of RARα403, thus resulting in less BarHL1^+^ cells inside the RP. P) Quantification of the total number of BarHL1^+^ cells displaying a mild reduction upon treatment with Su(H)*^DBM^*. N = 9,10,9,8 embryos for control, RARα403, RARα403+Su(H)*^DBM^*and control+Su(H)*^DBM^* groups, respectively. *p < 0.05, ***p < 0.001, Welch’s t test for A-J, Two-way ANOVA test for O-P. Scale bar, 50 μm.

#### Segregation of RP from dI1 interneurons requires RA inhibition of Notch signaling

Whereas the temporal separation between NC and RP traits depends on repression of BMP/Wnt activities by RA^15^, the spatial segregation between RP and adjacent dI1 interneurons is less well understood. Ofek et al. demonstrated that Notch signaling is responsible for the formation of both RP and dI1 interneurons, likely by defining a boundary between both cell types, while bearing no earlier effect on NC emigration ^16^. Reciprocally, in absence of RA activity, dI1 neurons infiltrate into the RP domain, suggesting that the boundary between them is compromised ^15^. Examination of the differentially expressed genes in response to RARα403 in the RP, revealed an upregulation of Notch-related genes (Fig. 2A), implying a potential repression of the Notch pathway by RA. Next, ISH of two central Notch pathway components – *DLL1* and *LFNG*, confirmed that their expression broadened into the RP domain of the RARα403-treated embryos. This stands in contrast to controls where the dorsal limit of gene expression corresponded to the boundary between RP and dI1 neurons (Fig. 5A-F and also ^16^).

Subsequently, we visualized Notch activity in the absence of RA signaling, implementing the pTP1 unstable reporter. Fig. 5G-J shows that control embryos displayed a narrow domain of Notch activity at the RP-dI1 boundary, while inhibition of RA activity led to the expansion of Notch activity throughout the entire RP. Together, these results indicate that RA represses Notch activity in the RP, confining it to boundary regions at this stage.

Thus, we reasoned that suppression of Notch signaling in cells deprived on RA activity, would rescue the boundary between RP and BarHL1-positive dI1 interneurons. To this end, we locally inhibited the Notch pathway in the dorsal NT by electroporating a Su(H) mutated in the DNA-binding domain (Su(H)*^DBM^*), expressed under the regulation of a RP-specific *ALDH1A2* enhancer. Using this construct, combined with inhibition of RA activity via RARα403, we reveal that while RA inhibition alone resulted in the abnormal presence of multiple BarHL1^+^ cells in the treated RP, this effect was significantly diminished when simultaneously inhibiting Notch activity (Fig. 5K-O). As anticipated, Su(H)*^DBM^*alone exhibited no discernible effect on dI1 cell distribution compared to control embryos, since Notch activity is typically absent in the RP itself (Fig.5O). A mild decrease in the overall count of BarHL1^+^ cells was monitored in embryos treated with Su(H)*^DBM^*, in both control and RARα403 conditions (Fig. 5P), consistent with the requirement of Notch activity for the generation of dI1 interneurons. Yet, the main effect, differentiating between controls with or without Su(H)*^DBM^*and RARα403-treated embryos with or without Su(H)*^DBM^*, was the spatial distribution of interneurons.

Together, the precedent results suggest that by repressing Notch activity in the RP and restricting it to the RP-dI1 interphase, RA indirectly defines the boundary between RP and dI1 domains.

### RA signaling induces fate segregation of late emerging NC-derived cell types

#### Gene expression profiles in control and RARα403-treated NC derivatives

The last NC cells to exit the trunk-level NT give rise sequentially to neural progenitors that settle in the periphery of DRG, and to melanocytes ^8,39^. Because of the technique utilized to isolate NTs, certain NC-derived cells expressing Msx1-Citrine and located in the mesenchyme near the NT were incorporated into our scRNA-seq analysis as clusters 11, 19, and 20 (Fig. 1E, H). Marker gene expression in the control sample revealed three distinct populations within the peripheral clusters: melanocytes (cl. 20), DRG progenitors (pDRG, cl. 11) and differentiating DRG neurons (cl. 19) (Suppl. Fig.6A-C). Apart from cell type-specific genes, DRG progenitors uniquely expressed *EDNRB,* whereas melanocytes revealed the presence of *EDNRB2*, two guidance receptors responsible for ventral vs. dorso-lateral routes of NC migration, respectively (Supp. Fig.6D) ^40,41^. The differentiating DRG cluster was characterized by expression of specific sensory neuron genes such as *NEUROD1, ISL1* and *BRN3*, as well as by generic neuronal genes like *DCX, NEFL,* and *TAGLN3* (Suppl. Fig.6A-C).

**Figure 6.**
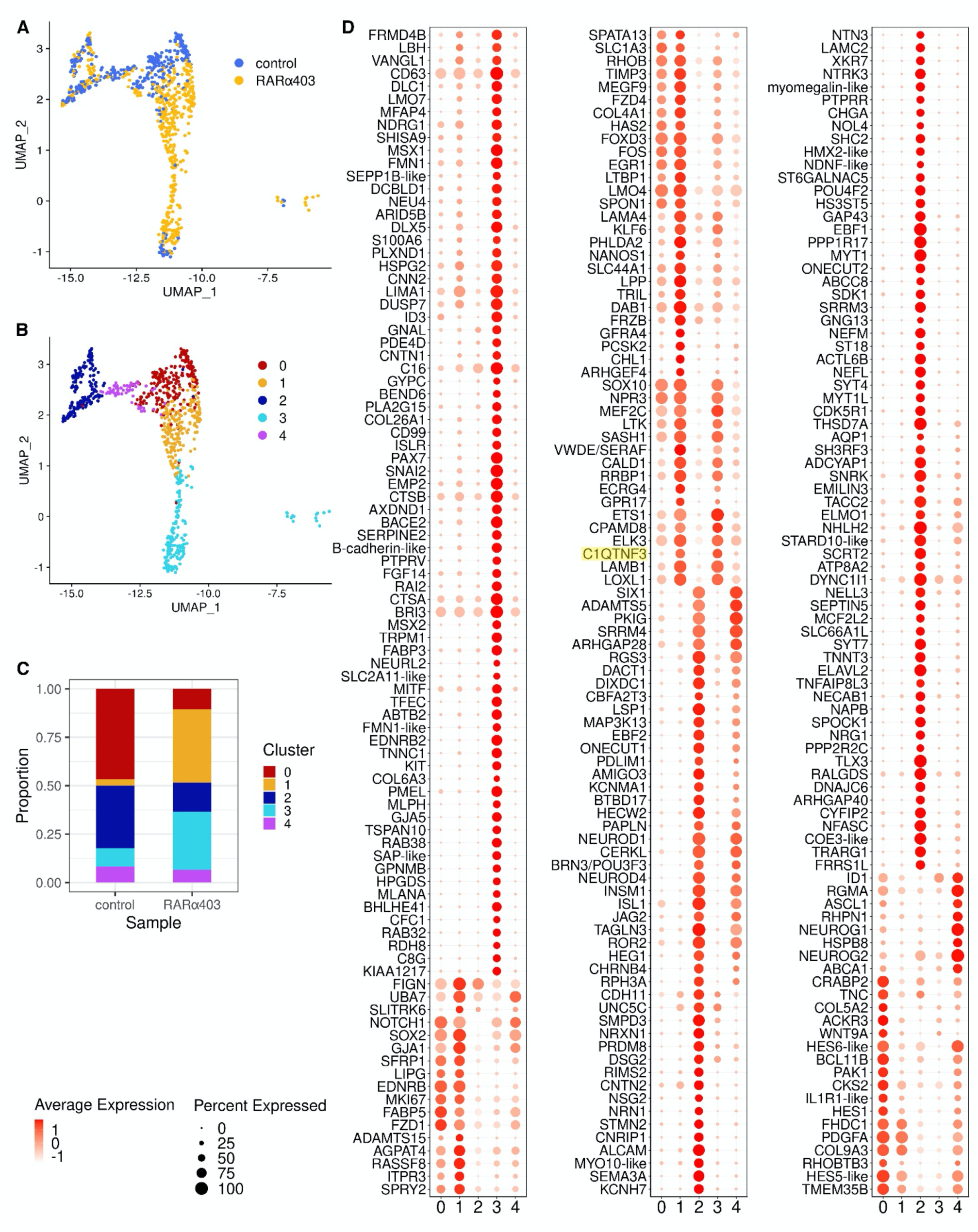
Re-clustering of NC derivatives reveals a bridge connecting DRG and melanocyte progenitors. A) UMAP visualization of the peripheral clusters (11, DRG progenitors; 19, differentiating DRG neurons; and 20, melanocytes) colored by sample. See Fig. 1E,H for cluster identification. B) Re-clustering generated five subclusters (0-4). Cluster 1 bridges between DRG progenitors and melanocytes. C) Proportions of the different subclusters, compared between control and RARα403 samples. Note that cluster 1 is predominant in the treated sample. D) Dot plots illustrating gene expression in the five subclusters, comprising both percentage of expressing cells and average expression level. Marker analysis was conducted on the data from peripheral clusters only. Highlighted in yellow is *C1QTNF3*, which is discussed below.

By comparing RARα403-treated and control samples, several gene categories displayed significant changes (Suppl. Fig.7). Notably, these effects were prominent in the pDRG and melanocyte clusters, while minimal differential gene expression was observed in differentiating DRG neurons. Importantly, a significant upregulation of glial genes such as *MBP, PMP22, KNDC1,* and of *VWDE/SERAF* ^42,43^, was monitored not only in the treated pDRG cluster, but also in the melanocyte group. In addition, *SOX2* was upregulated in pDRG, suggesting the maintenance of a progenitor state in this cell cluster.

**Figure 7.**
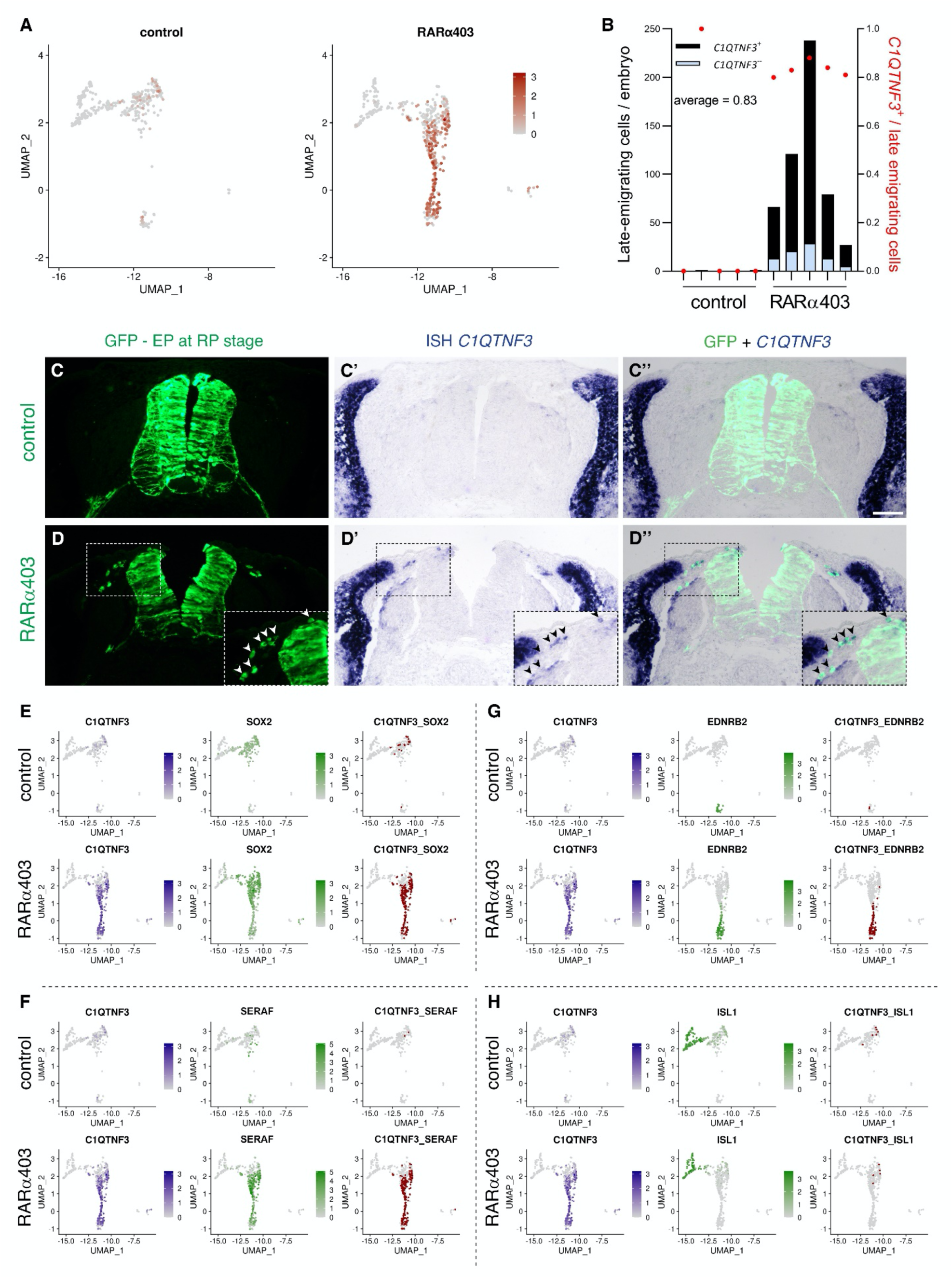
The bridge connecting DRG and melanocytes corresponds to late-emigrating cells and is defined by C1QTNF3 expression. A) UMAPs depicting expression of *C1QTNF3* in the peripheral clusters of control and RARα403 samples. Expression is primarily restricted to cells in the treated sample that connect between DRG progenitors and melanocytes. B-D”) Expression of *C1QTNF3* in late-emigrating cells. Embryos were electroporated at E2.5 with either control PCAGG or RARα403, and NTs were subjected to a second electroporation with a GFP plasmid at E3.5 to label the behavior of late NC cells. Embryos were fixed at E4.5 and GFP^+^ late-emigrating cells were in situ hybridized for expression of *C1QTNF3.* B) Quantification of the number of late-emigrating cells per embryo with and without expression of *C1QTNF3*. C-D”) Whereas virtually no positive cells exited the control NTs, numerous NC cells left the NT by E4 in the treated samples. Note in D-D’’ (inset, arrowheads) the presence of emigrated cells co-expressing GFP and *C1QTNF3*. N = 5,5 embryos for control and RARα403. E-H) Co-expression of *C1QTNF3* with other genes in both control and RARα403 samples visualized on UMAPs of the peripheral clusters. Double-positive cells are marked in red in the right columns. (E-G) Note the negligible proportion of double-positive cells in the control sample, as opposed to significant co-expression of *C1QTNF3* with progenitor (*SOX2*), glia (*SERAF*) and melanocyte (*EDNRB2*) genes in the bridge of the treated sample. H) No co-expression is detected between *C1QTNF3* and the neuronal gene *ISL1*. Additional visualizations are depicted in Supplem. Fig. 8 and see Fig. 9A for quantification.

An increase in the expression of ECM-related genes (such as *COL2A1, LAMB1, LAMA4*) and genes associated with adhesion, EMT and migration (such as *DUSP7, OLFM1, MEF2C*) was also observed in the treated pDRG and melanocyte clusters (Suppl. Fig.7). This suggests that the absence of RA activity encourages cellular remodeling.

#### A bridge connecting DRG progenitors with melanocytes corresponds to the late-emigrating

##### NC cells observed in RARα403-treated embryos

Whereas pDRG and melanocyte clusters were separated in the control UMAP, a ’bridge’ connecting these clusters emerged in the RARα403 sample (Fig. 6A). To better characterize it, the three peripheral groups were re-clustered, resulting in five distinct subpopulations (Fig. 6B-C) with unique gene expression profiles (Fig. 6D). The bridge was primarily identified as cluster 1 while also containing part of cluster 3. Cluster 1 was prominent in the treated sample but nearly absent in the control (Fig.6A-C). Examination of marker genes unveiled that cluster 1 is characterized by a range of glia-related genes, including *EDNRB, FOXD3, SERAF, SOX2, GPR17*, etc. Some markers of cluster 1 were also shared with cluster 3 (originally the melanocytic cluster). These comprised *SOX10, SASH1, CALD1, NPR3, MEF2C, LOXL1*, etc. Additionally, it is worth mentioning *DLC1*, recently found to mediate cranial NC delamination and migration ^44^, and *CCN2*, a glial ECM protein with EMT properties ^45^. Thus, emergence of the bridge cluster in the treated sample confirms the maintenance of a link between DRG progenitors and melanocytes in absence of RA activity.

We previously reported that inhibition of RA signaling in the dorsal NT prior to NC-to-RP transition, extends the period of NC production and emigration into the RP stage, when no NC cells are normally produced any longer ^15^. Since these late-emigrating cells were unique to RARα403-electroporated embryos, we set to examine whether bridge cells, also specific to the treated embryos, correspond to the late-emigrating cells observed in vivo. The most distinctive marker of the bridge cells was the adipokine *C1QTNF3* ^46^, a factor with unknown function in the present context. *C1QTNF3* was highly specific to bridge cells and absent from additional cell subsets in our dataset (Figs. 6D and 7A). ISH confirmed *C1QTNF3* expression in 83% of the late-emigrating cells, yet no expression was detected in neural cells of controls (Fig. 7B-D”). Furthermore, co-expression analysis of *C1QTNF3* with an array of genes revealed that a significant proportion of bridge cells co-expressed *SOX2*, further confirming their persistent progenitor state. Additionally, they co-expressed an assortment of glial (e.g. *SERAF, MBP, KNDC1, EDNRB*) and melanocytic genes (e.g. *EDNRB2, PMEL, TFEC*) but not of neuronal genes (e.g. *ISL1, NEUROD1, NTRK3*) (Fig.7E-H, Supplem. Fig.8).

**Figure 8.**
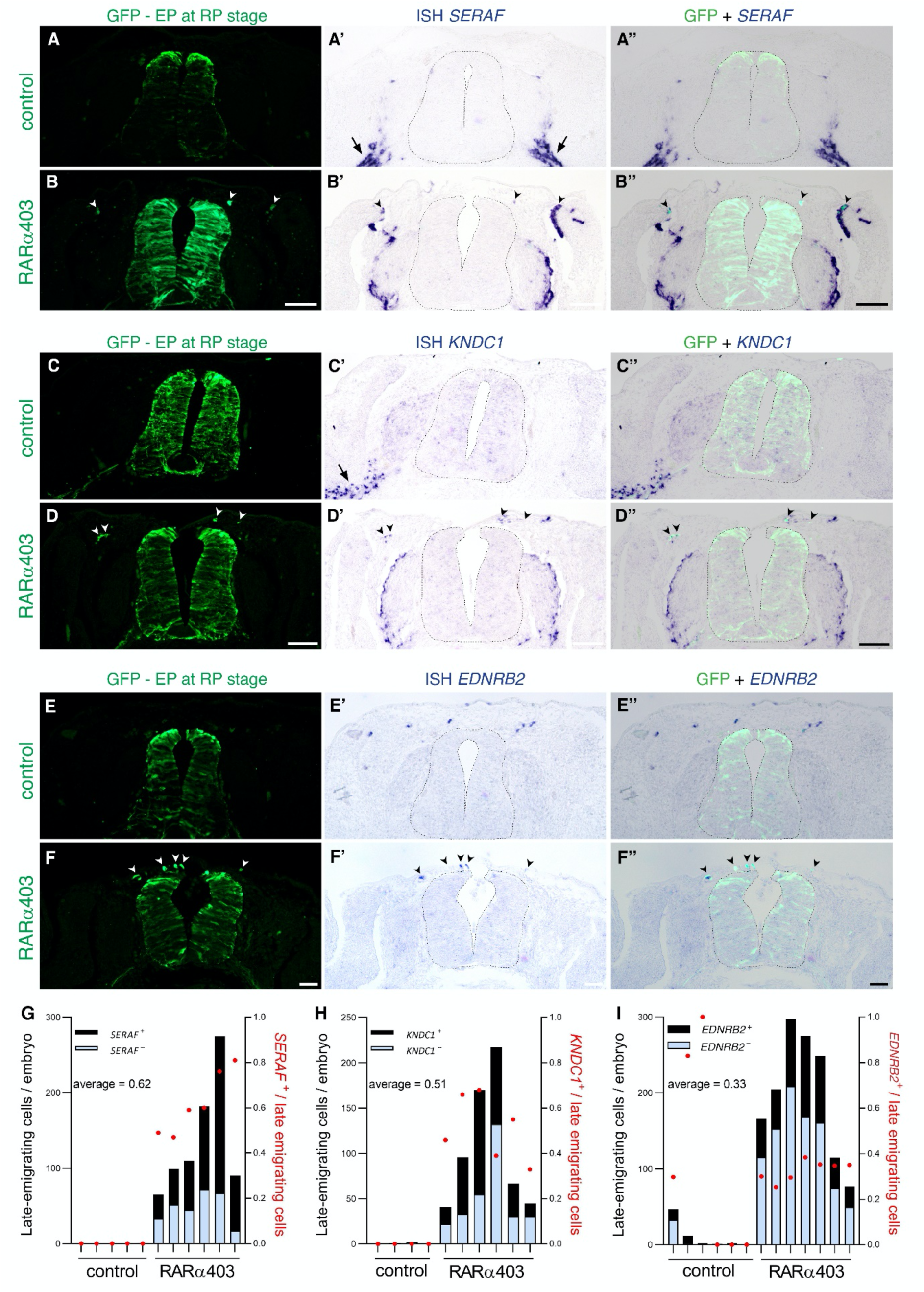
The late-emigrating bridge cells express glial and melanocyte markers. In vivo validation of the expression of the glial genes *SERAF* and *KNDC1* (A-D, G-H) and the melanocyte marker *EDNRB2* (E-F, I) in late-emigrating cells. Embryos were electroporated at E2.5 with either control PCAGG or RARα403, and NTs were labeled with GFP via a second electroporation at E3.5. Embryos were fixed at E4.5 and GFP^+^ late-emigrating cells co-expressing the various genes were counted. A-D) ISH for *SERAF* (A-B) and *KNDC1* (C-D). Arrows indicate normal expression in progenitors residing along nerves, and arrowheads mark GFP^+^ late-emigrating cells in DRG and throughout the mesoderm restricted to the treated samples (see Supplem. Fig.9 for additional images). E-F) ISH for *EDNRB2*. Note expression in control melanocytes (E-E’), and co-expression with GFP in late-emigrating cells (F-F’’). G-I) Quantification of the expression of *SERAF*, *KNDC1* and *EDNRB2* by late-emigrating cells. N=5,6 for *SERAF*; N = 4,6 for *KNDC1* ; N = 6,7 for *EDNRB2,* in control and RARα403, respectively.

Moreover, ISH for the glia-specific genes *SERAF* and *KNDC1*, showed their expression in Schwann cells residing along the spinal nerve of control embryos (Fig. 8A-A’, C-C’ and Supplem. Fig9A,D, arrows). In treated embryos, they were additionally apparent in the DRG periphery, in the mesenchyme surrounding the NT and also in dermis, where melanocytes typically migrate (Fig.B-B’, D-D” and Supplem Fig.9B,C,E,F, arrowheads). Specifically, *SERAF* and *KNDC1* were expressed by 62% and 51% of late-emigrating cells, respectively (Fig.8G,H). In addition, EDNRB2, transcribed by melanocytes of control embryos (Fig8E-E”), was also detected in 33% of late-emigrating cells and was mainly detected throughout the dermis (Fig.8F-F”, I). Collectively, these findings confirm that late-emigrating cells express glia and melanocytic genes, thus providing the in vivo equivalent of the bridge cells connecting pDRG and melanocytic clusters in the scRNA-seq.

**Figure 9.**
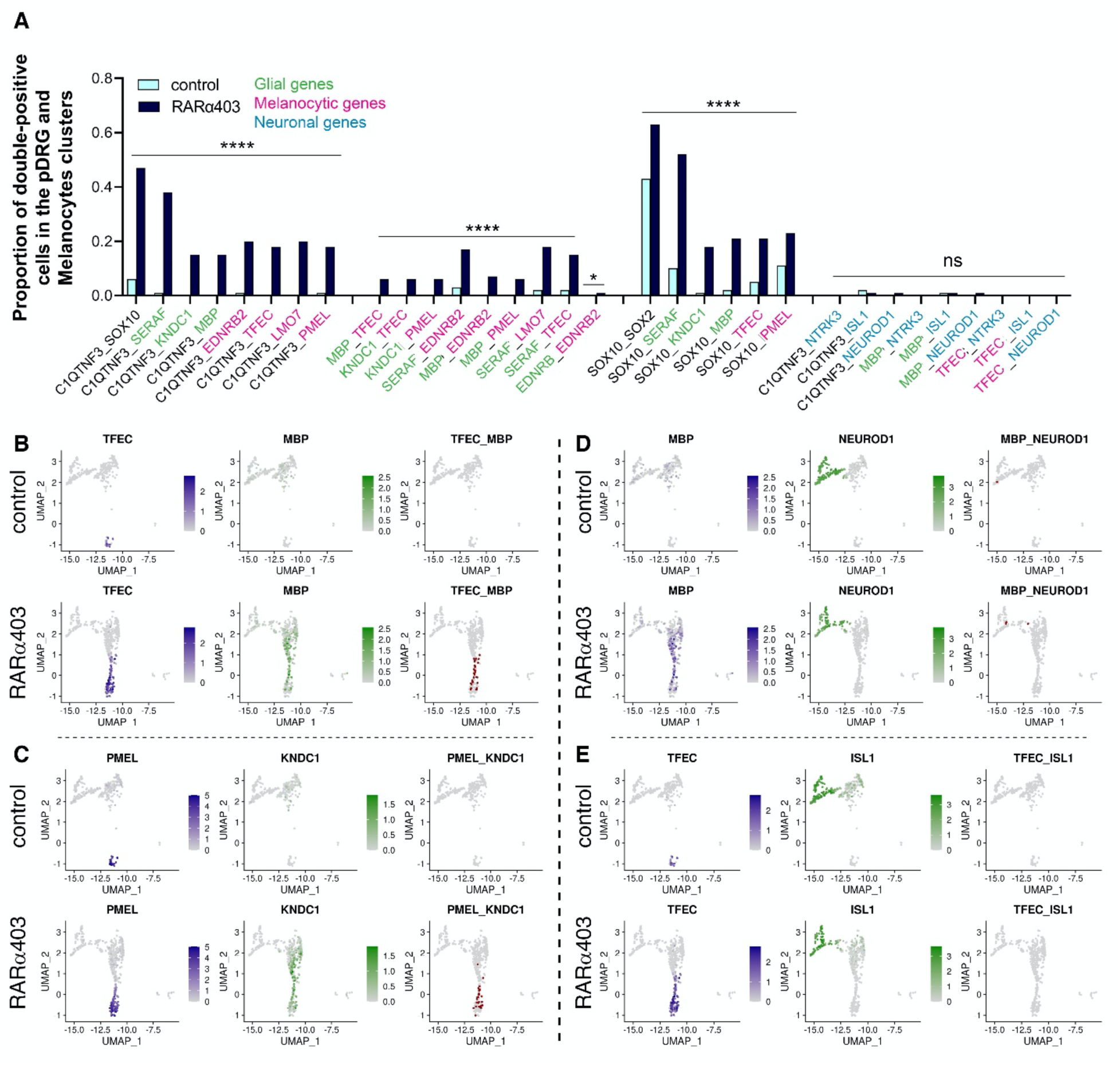
Single cells in the bridge co-express glia and melanocyte genes. A) Testing for the co-expression of an array of gene pairs in pDRG and melanocyte clusters (see Methods). From left to right, *C1QTNF3* combined with glia or melanocyte genes, glia combined with melanocyte genes, *SOX10* combined with glia and melanocyte genes, and neuronal genes with *C1QTNF3* and glia/melanocyte markers. B-E) Visualization of selected pairs from the above quantifications on UMAPs of the peripheral clusters, separated into control and RARα403 samples. *p < 0.05, ****p < 0.0001

#### A common glia-melanocyte (GM) progenitor is revealed in absence of RA signaling

The precedent results suggest that bridge cells might represent a transitional state preceding commitment to either glial or melanocytic fates. This would predict that individual glial progenitors co-express melanocyte traits. In fact, the existence of a bipotent NC-derived GM progenitor was proposed based on in vitro studies^17^. Moreover, Schwann cell progenitors were also demonstrated to generate melanocytes in vivo ^47–49^see Discussion). Thus, we examined co-expression of glia, melanocyte, and neuronal genes in single cells within the pDRG and melanocyte clusters. Co-expression of glia with melanocyte genes was evident in a subset of the RARα403-treated sample (Fig.9A-C, Supplem. Fig.10 A-E), while being nearly absent in controls. In contrast, neuronal genes were restricted to the differentiating DRG cluster, showing no co-expression with either glial or melanocyte genes (Fig. 9A, E, and Supplem. Fig. 10F). *SOX10* is normally transcribed in both early glia and pigment lineages ^50,51^. Consistently, it was present in almost all cells within the pDRG and melanocyte clusters of control samples, serving as a positive control in the co-expression analysis. Furthermore, an enhanced proportion of *SOX10*+ cells combined with either glia or melanocyte genes was monitored upon treatment (Fig. 9A). Hence, our bioinformatic data suggest that lack of RA signaling supports the maintenance of a common GM progenitor while preventing its timely segregation into individual fates.

**Figure 10.**
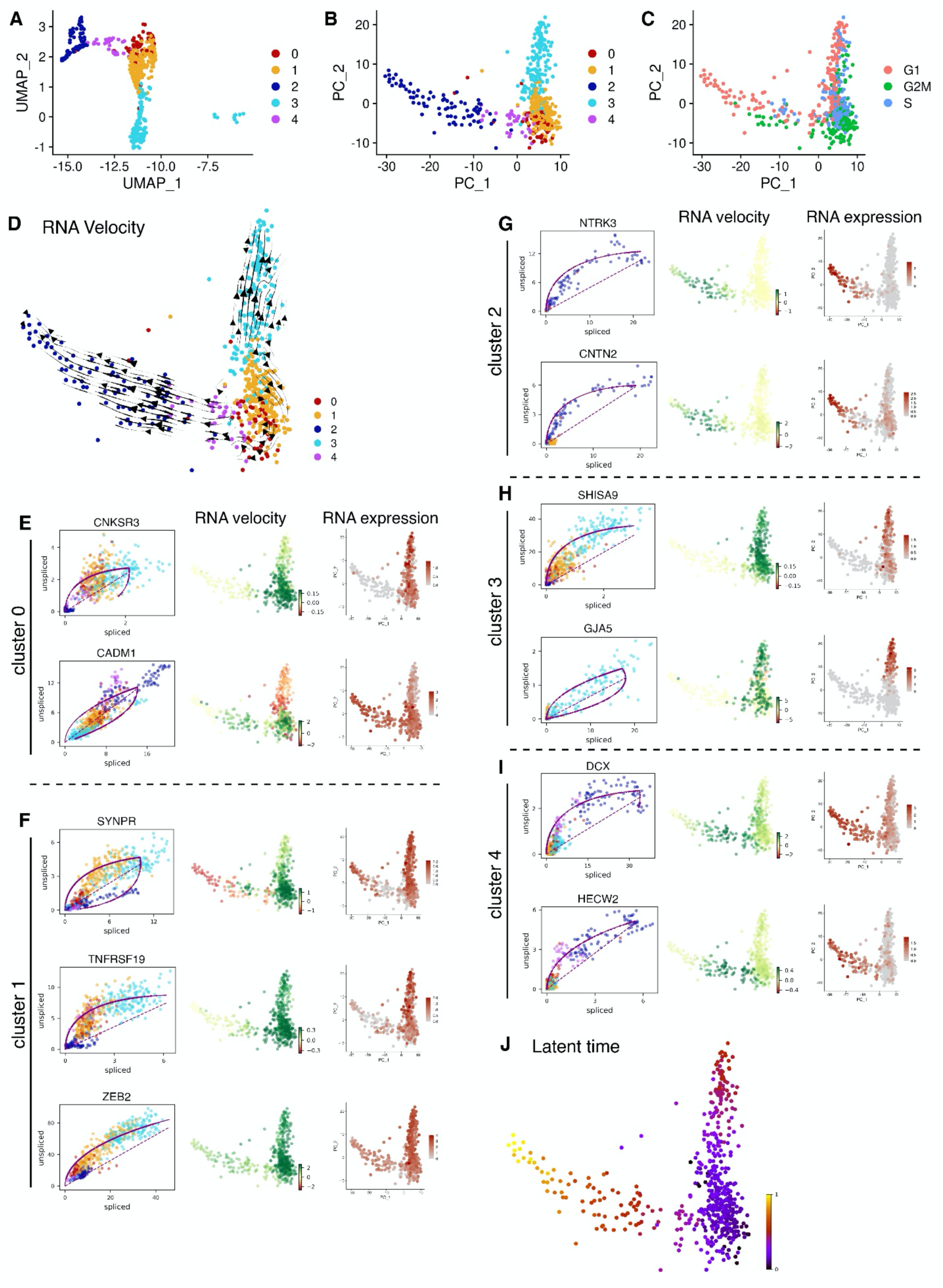
Velocity analysis reveals bridge cells as an origin for melanocytes and DRG cells. A-B) Re-clustering of the peripheral cell types projected on UMAP and PCA, respectively. C) PCA colored by the cell cycle phase (predicted using Seurat’s CellCycleScoring function). D) RNA velocity vectors projected on a PCA of peripheral clusters. E-I) Examples of specific genes with high velocity in different subclusters. Left, phase portraits of spliced versus unspliced transcripts, with a dotted line indicating the steady state of transcription and a fitted curve indicating the learned dynamics; middle, PCA colored by the velocity; right, RNA expression level. J) Latent time analysis of peripheral clusters. Earliest time point is depicted in black/purple and latest in yellow.

Next, to obtain a dynamical view of the above findings, we estimated RNA velocities of the treated sample using the dynamical model implemented in scVelo, an unbiased approach that leverages the distinction of unspliced and spliced RNA transcripts from the aligned sequences, enabling to infer whether the expression of a gene is being initiated or downregulated, respectively^52^. For improved visualization, a PCA projection instead of a UMAP was applied (Fig. 10A-B). Cell cycle analysis suggested that the separation into clusters is unlikely to be affected by the cell cycle phase (Fig. 10C). In our dataset, RNA velocity revealed a clear bifurcation stemming from cluster 1, the bridge, leading either towards melanocytes (Cluster 3) or towards DRG progenitors (Cluster 0) followed by neuronal progenies (Clusters 4 and 2) (Fig. 10D). Next, we interrogated the behavior of single genes in each cluster. For example, cluster 0 (pDRG) top velocity genes such as *CNKSR3* and *CADM1*, stabilized towards either melanocytes or neurons, respectively (Fig. 10E). Cluster 1 genes with highest velocity in bridge cells, streamed towards pDRG and melanocytes (Fig. 10F, Supplem. Fig 11A). Furthermore, Cluster 3 genes streamed towards a melanocyte fate yet were also upregulated in bridge cells (Fig. 10H, Supplem. Fig.11C). Thus, the dynamic trajectories leading from bridge progenitors towards segregated fates are effectively illustrated by analysing single genes. In contrast, genes with top velocity in clusters 2 or 4 were mainly expressed in differentiated or differentiating neurons, respectively, consistent with their patterns of mRNA expression (Fig. 10G,I, Supplem. Fig. 11B,D). Finally, scVelo’s latent time analysis further confirmed that pDRG and bridge cells are produced earlier than melanocytes and neuronal cells (Fig. 10J).

## Discussion

By implementing a combination of *in-ovo* gene misexpression and single-cell RNA profiling, we here demonstrate that RA is an essential component of the nascent RP organizer, that plays a key role in segregation of cell fates in both time and space.

We have recently reported that RA is responsible for the end of NC production and emigration hence segregating between PNS (NC) and dorsal CNS (RP) fates. This is accounted for by repressing BMP and Wnt activities ^15^, both found to be necessary for the onset of NC EMT ^4,53^. Consistent with these findings, our scRNA-seq analysis now confirms the RA-dependent upregulation of specific BMP inhibitors in RP, that were previously uncovered in a differential transcriptome analysis between premigratory NC and RP ^15,16^. These repressors are likely to mediate, at least partially, the loss of responsiveness to BMP, as documented for early misexpression of Hes1 ^9^.

In the absence of local RA activity, not only NC cells fail to separate from the RP lineage in the temporal dimension, as we also observe an invasion of the RP by dI1 interneurons, but not by more ventral interneuron types. This suggests that, even if a RP forms without RA, it fails to spatially separate from its ventrally localized neighbours. Notably, previous results documented that Notch signaling is both necessary and sufficient for the formation of a RP and of dI1 interneurons ^16^, and here we further uncover that separation of RP from dI1 neurons is elicited by RA via restriction of Notch signaling to the boundary region between both cell types. This is consistent with loss of the constitutive expression of the boundary gene Hes1/4 ^54,55,56^ in the treated RP, and may also be related to the observed downregulation of *ISM1*, a nodal repressor expressed in the midbrain-hindbrain boundary ^55,56^. Associated with the compromised separation of RP from dI1 lineages, absence of RA input might lower the mechanical tension required for keeping the boundary straight and precluding cell intermixing^57^.

Our results further uncovered changes in various axonal guidance genes, and consistent with these observations, the growth of dI1 axons was aberrant and extended outside the confines of the NT. Which of the pathways modified by lack of RA (*SLIT1, CXCR4, EPHA4*, etc) accounts for the phenotype observed, remains to be clarified. Together, our results illustrate that RA is used reiteratively and contextually in combination with BMP, Wnt and Notch, providing further insight into how a small repertoire of signaling pathways interacts to ensure that correct cell types are specified at the right time and place. In addition, despite being consistently produced in the paraxial mesoderm ^58^, the RA responsible for the regulatory functions here defined originates from a novel source — the nascent RP ^15^. This emphasizes the concept that different sources of the factor are associated with distinct functions ^59–61^.

Along this line, loss of RP-derived RA activity is associated with substantial cellular alterations. Treated RPs are loosely organized, as evidenced by loss of cell adhesion molecules, upregulation of metalloproteases, maintenance of a discontinuous basal lamina and upregulation of EMT factors, when compared to the normal RP that is an epithelial structure with distinct boundaries (this study and see also ^15^). A highlight of this plasticity is an altered state of cellular commitment, as the RP of RARα403-treated embryos remains proliferative ^15^, and exhibits the presence of single progenitors co-expressing NC, RP, and dI1 markers. While the lack of segregation between NC and RP lineages is less surprising due to their shared derivation from a FOXD3^+^ lineage ^8^, the presence of a common progenitor for NC/RP and dI1 cells is more intriguing and suggests that boundaries between domains serve not only structural purposes but also facilitate the correct segregation into distinct cell types. For instance, in controls, dI1 progenitors exhibit high BMP activity as opposed to their RP neighbours. Perhaps the extension of BMP activity to the RP of RARα403-treated embryos ^15^, coupled with aberrant Notch activity, enables dI1 features to be expressed inside the RP domain and prevents phenotypic segregation.

Mechanical harvesting of the NTs for RNA-seq purposes, also enabled the collection of Citrine^+^ NC-derived cells encountered in the mesoderm adjacent to the NT. These cells were also likely to be exposed to dorsal NT-derived RA, as its synthesis begins at the late NC stage in dorsal NT, just prior to the establishment of the definitive RP ^16^. Bioinformatic analysis revealed that, in addition to distinct clusters composed of DRG progenitors, DRG neurons or melanocytes, all apparent under control conditions, an additional cluster appeared only in the absence of RA activity. We termed this cell group a “bridge” as it connects DRG progenitors with melanocytes. Molecularly, bridge cells are composed of both DRG progenitors with a glial character and/or also melanoblasts, but not neuroblasts or neurons. Most notably, GM traits were expressed by single cells of treated samples, yet few or no cells co-expressed these genes under control conditions. Hence, bridge cells represent a “hybrid” state along the hierarchy of cell specification, unravelling in vivo, the presence of a bi-fated intermediate cell type with both GM progenitor properties, and further substantiating the notion that RA is necessary for fate segregation between the above peripheral lineages.

Our findings are in line with previous studies highlighting the plasticity of the melanocyte-glial state in vitro by showing that these phenotypes can interconvert under specific medium conditions ^17,18,62^. In mice, the decision to become glia or melanocytes was shown to be driven by Wnt signaling that acts upstream of Sox10 and Pax3 ^20^, or by NR2F1 in a model of Waardenburg syndrome ^63^. Moreover, mice carrying spontaneous or targeted mutations of *Sox10* lack satellite glia and Schwann cells and exhibit pigmentation defects but neurogenesis is initially unaffected ^64^. Furthermore, melanoma progression from normal pigment cells is accompanied by a de-differentiation state in which melanocytic markers such as *DCT* and *MITF* are downregulated and glial/melanoblast markers are instead upregulated ^65^.

Most notably, Schwann cell progenitors, a specialized glial subset, generate either Schwann cells or hypaxial and limb melanocytes depending on the level of contact with peripheral nerves^48^. This process is regulated by cross-repressive interactions between FOXD3 and MITF ^49,66,67^.

In addition to melanocytes derived from Schwann cell progenitors, epaxial pigment cells have a direct origin from the NC following migration through a dorso-lateral pathway ^49^. Our RNA-seq analysis supports the presence of two distinct pigment cell populations, one with melanocytic genes being exclusively expressed in the melanocyte cluster and another shared with bridge and presumptive glia cells. This raises the interesting hypothesis that bridge cells might represent a population of progenitors able to generate either ganglionic glia, Schwann cells or melanocytes. This is consistent with the maintenance in bridge cells of *FOXD3*, *SOX2* and *SOX10*, markers of the precursor state. RNA velocity and latent time analyses support the notion that bridge cells are early progenitors from which a bifurcation into either melanocytes or neural progenitors evolves.

In the current study, we have also delineated the bridge subpopulation, as corresponding to late-emigrating NC progenitors that become apparent only in the absence of RA signaling ^15^. A noteworthy finding is that over 80% of these late-emigrating cells express a distinctive gene, which is novel in the present context — the adipokine C1QTNF3 (CTRP3). C1QTNF3, known as an anti-inflammatory cytokine ^46,68,69^, plays a role in stimulating macrophage chemotaxis during adipose tissue remodelling ^70^. Intriguingly, it is one of the 12 prognostic genes linked to recurrence-free survival in human cutaneous melanoma ^71^ and is a significant gene associated with pigmentary traits ^72^. Our analysis has revealed that late-emigrating, *C1QTNF3*^+^ bridge cells co-express both glial and melanocyte genes, excluding neuronal genes. This suggests a potential involvement in the development of glia-melanocyte interactions, warranting further exploration into potential developmental functions.

Another intriguing gene in the present context is *SERAF*. *SERAF* was initially recognized in avian embryos as an early, Schwann cell-specific gene regulated by SOX10 ^42,43^. In control embryos at E4, *SERAF* showed predominant expression in Schwann cell progenitors along peripheral nerves. However, in treated embryos, it marked over 60% of late-emigrating cells and distributed throughout the mesenchyme, including subectodermal sites and the periphery of DRG, a region where satellite glia develops. This suggests that endogenous RA signaling may restrict *SERAF* expression to the Schwann cell lineage, and the absence of RA allows for the emergence of a multipotent progenitor that includes satellite glia. In contrast, bridge cells are unlikely to represent boundary cap cells ^73,74^, as no *KROX20 (EGR2)* was detected, and the expression of *PRSS56*, another specific marker^75^, remained unchanged in the absence of RA activity.

Collectively, our data not only shed light on the lineage potential of RA-deficient cells but it may also infer on aspects of normal fate segregation influenced by endogenous RA activity. In this context, it is tempting to suggest the existence of a common NC/RP/dI1 progenitor at some point in early neural tube development. Moreover, our findings hint at the possibility that at least a subset of glia and melanocytes evolves from a shared GM progenitor during normal development.

## Materials and Methods

### Embryos

Fertilized quail (*Coturnix coturnix Japonica*) eggs were obtained from commercial sources (Moshav Mata), kept at 15 °C and then incubated at 38 °C to desired stages. Embryos were staged by the number of somite pairs formed. All experiments were conducted at the flank level of the axis.

### Expression vectors and in-ovo electroporation

The following expression vectors were used: control pCAGG, pCAGGS-eGFP ^8^, pCAGGS-RFP ^16^, pCAGGS-RARα403 ^15^, Msx1(ehn-264)-Citrine ^21^, Atoh1-tdTomato-F, Atoh1-GFP ^76^, aldh1a2-1G-eGFP ^77^, aldh1a2-1G-RFP (subcloned from Castillo et al, 2010, eGFP replaced with RFP), aldh1a2-1G-xSu(H)1 *^DBM^* (subcloned from Castillo et al, 2010, replaced with xSu(H)1 *^DBM^* ^78^, pCA-FoxD3-IRES-EGFP^79^, pCAGGS-cSnai2(Slug)-IRES-nls-GFP, pCAGGS-Sox9-IRES-nls-GFP ^80^, and pTP1-d4Venus ^81^

For NT electroporation, DNA (4 ug/ul) was mixed with Fast Green and microinjected into the lumen of the NT at the flank level of the axis. Five mm tungsten electrodes were placed on either side of the embryo. A square wave electroporator (BTX, San Diego, CA, USA) was used to deliver one pulse of current at 17–25 V for 6ms.

### NT collection and dissociation

Embryos were electroporated at E2.5 with either control or RARα403 vectors, together with a Msx1-Citrine plasmid. At E4, ten NTs per group (with associated mesoderm) were mechanically microdissected in Phosphate-buffered saline (PBS) supplemented with Ca_2_^+^/ Mg_2_^+^ and 5% fetal calf serum (FCS). NTs were incubated for 15 min at 37°C in 1.5 ml Accutase (Sigma Israel, A6964) containing 30 ul Papain (Sigma Israel, P3125) and 300 ng DNaseI (Sigma Israel, DN25) and dissociated mechanically using sequentially 1000 and 200 ul pipettes. Cells were filtered through a 40-micron filter and resuspended in PBS containing 2% FCS. Samples were sorted using a BD FACSAria III sorter (BD Biosciences UK). Single cells were isolated by sequentially gating cells according to their SSC-A vs. FSC-A and FSC-H vs FSC-W profiles, following standard flow cytometry practices. Dead or damaged cells were excluded following Propidium Iodide uptake. Citrine fluorescence was detected using a 488 nm laser; age-matched NT cells from untransfected embryos served as negative control to determine the fluorescence gate.

### Single-cell RNA sequencing and analysis

Sorted fluorescent cells were loaded into the 10X Genomics Chromium Next controller. Libraries were prepared following manufacturer’s instructions (GEM Single Cell 3ʹ GEM, Library & Gel Bead Kit v3.1, 10× Genomics, CA, USA). Approximately 10.000 cells were loaded per sample. Sequencing was performed using the Illumina Nextseq500 platform (Illumina Inc, San Diego, CA,USA) with the following sequencing conditions: 28 bp (Read1) and 54 bp (Read2).

The Cell Ranger pipeline ^82^ (v6.0.1, 10x Genomics) with default parameters was used for alignment, filtering, barcode counting and Unique Molecular Identifier (UMI) counting. The Coturnix coturnix japonica 2.1 genome sequence and annotations were downloaded from NCBI (RefSeq assembly GCF_001577835.2). The gtf was filtered using Cell Ranger’s mkgtf command to keep protein-coding genes and lncRNAs.The Seurat R package ^83^ (v4.04) was used for downstream analysis and visualization. Gene-cell matrices were filtered to remove: (1) cells with mitochondrial reads comprising more than 8.5% of all reads; (2) cells with less than 250 genes or more than 7500 genes; (3) cells that had less than 500 UMIs or more than 40,000 UMIs. In addition, genes detected in fewer than 10 cells were excluded from the analysis. After implementing these quality control measures, a total of 6,164 control cells, 10,191 RARα403 cells, and 16,706 genes were retained. A cluster consisting of low-quality cells (1,789 cells that had a low number of detected genes and low number of UMIs) was excluded. After that filtration, 5,890 control cells and 8,676 RARα403 cells were retained for further analysis.

The expression data was normalized and log-transformed using Seurat’s NormalizeData function. The top 2,000 highly variable genes were identified using Seurat’s FindVariableFeatures function with the ‘vst’ method. Each cell was assigned a cell-cycle score using Seurat’s CellCycleScoring function and the NCBI’s orthologs of the G2/M and S phase marker lists from the Seurat package. Potential sources of unspecific variation in the data were removed by regressing out the mitochondrial gene proportion, UMI count and the cell cycle effect using linear models and finally by scaling and centering the residuals as implemented in the function “ScaleData” of the Seurat package. Principal component analysis (PCA) was performed. We selected 24 principal components (PC) for downstream analyses. Cell clusters were generated using Seurat’s unsupervised graph-based clustering functions “FindNeighbors” and “FindClusters” (resolution = 0.5). UMAP was generated using the RunUMAP on the projected principal component (PC) space. Seurat’s functions FeaturePlot and DimPlot were used for visualization. Seurtat’s DotPlot function was used to generate dot plots to visualize gene expression for each cluster. Plots were further formatted using custom R scripts with the packages ggplot2 and ggnewscale. R package version 0.4.8, and patchwork (Pedersen T, 2022, patchwork: The Composer of Plots. R package version 1.1.2, https://CRAN.R-project.org/package=patchwork). Heatmaps were produced with The R package ComplexHeatmap (Version 2.14) ^84^.

Marker genes for each cluster were identified by performing differential expression between a distinct cell cluster and the cells of the other clusters with the non-parametric Wilcoxon rank sum test (Seurat’s FindAllMarkers function). Only the control sample was used for the marker identification. Cell types were assigned manually based on the expression of classic marker genes.

In order to get higher clustering resolution, the RP cells (cluster 18) and the DRG-melanocyte cells (clusters 11, 19, and 20) were each re-analyzed with the workflow described above using the 500 most highly variable genes with the following parameters: (a) 20 PCs and resolution of 0.8 for the RP cells (b) 19 PCs and resolution of 0.4 for the DRG melanocyte cells.

### Testing for co-expression of marker pairs in RP and DRG-melanocytes

A marker was considered to be expressed in a cell if it had a count of at least 2. The number of cells that were positive for both markers was determined for each sample. A Chi-square test was used to test the null hypothesis that the proportions are equal (prop.test in R). P-values were corrected for multiple comparisons using the FDR method of Benjamini and Hochberg ^85^

Differential expression analysis inside clusters between conditions was performed using the non-parametric Wilcoxon rank sum test (Seurat’s FindMarkers function).

Out of the 16,706 genes in our data, 4422 had uncertain function (LOC symbols; LOC plus the GeneID). Of them, 40 LOC genes that exhibited notable expression patterns, were assigned names based on high similarity to orthologous genes (Supp. Table 1).

### RNA Velocity and Latent time analysis

Velocyto (version 0.17.17) ^86^ was used to generate count matrices, based on the spliced and unspliced reads of the RARα403-treated cells, using the run10x option. Unspliced and spliced counts were matched to the barcodes and genes retained after filtering. scVelo (version 0.2.5) ^52^was used to compute velocities using the following options: filter_and_normalize was run using min_shared_counts set to 20 and min_cells set to 80, monemts was run with n_neighbors set to 10. Velocities were calculated using the “dynamical” mode and visualized on the PCA. Phase portrait plots and PCA colored according to the velocity, were created using scVelo pl.velocity function. Latent time was calculated using scVelo tl.latent_time function. The RNA velocity analysis was performed without regressing out the cell cycle effect.

### Immunohistochemistry

For immunostaining, embryos were fixed overnight at 4 °C with 4% formaldehyde (PFA) in PBS (pH 7.4), embedded in paraffin wax and serially sectioned at 8 μm. Immunostaining was performed either on whole mounts or paraffin sections, as previously described ^87,88^. Antibodies were diluted in PBS containing 5% fetal bovine serum (Biological Industries Israel, 04-007-1A) and 1% or 0.1% Triton X-100 (Sigma Israel, X-100), respectively. Antibodies used were: rabbit anti-GFP (1:1000, Invitrogen, Thermo-Fisher Scientific, A-6455), chicken anti-GFP (1:500, Novus, NB100-1614), rabbit anti-Sox9 (1:150, Millipore, AB5535), rabbit anti-Snai2 (1:500, CST, CST9585), rabbit anti-BarHL1 (1:300, Sigma Israel, HPA004809), mouse anti-Neurofilament-associated (1:10, DSHB, 3A10), and mouse anti-Pax7 (1:10, DSHB, PAX7). Cell nuclei were visualized with 125 ng/ml Hoechst 33258 (Sigma Israel, 14530) diluted in PBS.

### In situ hybridization

For ISH, embryos were fixed in Fornoy (60% ethanol, 30% formaldehyde and 10% acetic acid) for 1 hr at room temperature, embedded in paraffin wax and serially sectioned at 10 μm. Briefly, sections were treated with 1 µg/ml proteinase K, re-fixed in 4% PFA, then hybridized overnight at 65 °C with digoxigenin-labeled RNA probes (Roche, 11277073910). The probes were detected with AP coupled with anti-digoxigenin Fab fragments (Roche, 11093274910). AP reaction was developed with 4-Nitro blue tetrazolium chloride (NBT, Roche, 11383213001) and 5-bromo-4-chloro-3’-indolyphosphate p-toluidine salt (BCIP, Sigma Israel, B8503). In the Snai2/*RSPO1* ISH experiment, the AP reaction of *RSPO1* was developed with Fast Red (Sigma Israel, F4648) for 2 hr at room temperature. Importantly, all hybridizations, whether done on intact embryos at NC vs. RP stages, or in control vs. experimental embryos, were always developed for the same time for a specific probe and experiment. The RNA probes used were produced either from a vector or using a PCR product using the KAPA2G Fast ReadyMix PCR kit, (Sigma Israel, KK5101). cDNA templates were synthesized by RNA precipitation followed by reverse transcription PCR. RNAs were produced from 20 somite-stage-to-E4 quail embryos. Tissue samples were homogenized with TriFast reagent, and RNA was separated with chloroform and isopropanol. RNA probes synthesized from plasmids were: *cBAMBI* ^89^, cEdnrb2 ^49^, *cLFNG* ^16^. Primers used for synthesizing probes via PCR were: *qDLX5* (GTTTGACAGAAGGGTCCCCA, CACTTTCTTTGGCTTGCCGT), *qKNDC1* (AGTTTGGCTGCGAATGGGAA, GCTCTAGGAAGGCTGGTACG), *qSERAF* (CCCTCAAGGTTGCAGGAATA, AACCGAGACATAGCAGTTCT), *qC1QTNF3* (AGTTTGGCTGCGAATGGGAA, GCTCTAGGAAGGCTGGTACG), *qNANOS1* (GCTTCATCACATTGCTCCAG, CTTCTGTGTTTGTAGCGAGC), *qRSPO1* ^15^, *qRSPO3* (TGCATCCTAACGTGAGCCAG, CACGGACTCCACTCACTAGC), *qADAMTS8* (TGATGGCCCCACTCTTTGTC, AGCCGTACTTGGATCGGTTG), *qALCAM* (GCGTCAAAACACGTAGACAA, CTTTAAGGCAGCGGATAACG), *qDLL1* (TACTGCACTGAGCCGATTTG, CACCATCAGGGTTGTCAGTG).

### Data acquisition and statistical analysis

Fluorescence intensity was quantified in 11-34 sections per embryo. The number of embryos monitored per treatment is depicted in the respective figure legends. Intensities of immunofluorescent and ISH signals were measured using FIJI ^90^. In most cases, the RP was selected as the region of interest (ROI); mean intensity was measured, and background intensity was then subtracted. Average intensity of all sections was calculated per embryo, and values were normalized with a control mean set as 1.

To monitor intensity of pTP1, a segmented line with an arbitrary thickness of 20 was drawn pursuing the curvature of the NT, with the dorsal midline in its center, and further measured along the line using FIJI. Subsequent analysis and graphics were performed using R (https://www.R-project.org/) in Rstudio (http://www.rstudio.com/). The code used and the source data are available on Github: https://github.com/dinarekler/pTP1.

The distance of BarHL1^+^ cells from the dorsal midline of the NT was measured for each cell manually using FIJI, and graphs were generated using R in Rstudio. The code used is available on Github: https://github.com/dinarekler/BarHL1-RARa403-SuHdbm.

Images were photographed using a DP73 (Olympus) cooled CCD digital camera mounted on a BX51 microscope (Olympus) with Uplan FL-N 20 x/0.5 and 40 x/0.75 dry objectives (Olympus). Confocal sections encompassing their entire thickness were photographed using a Nikon Eclipse 90i microscope with a Plan Apo 40 x/1.3 or 100 x/1.4 objectives (Nikon) and a D-Eclipse c1 confocal system (Nikon) at 1.0 μm increments through the z-axis. Images were z-stacked with FIJI software.

For quantification, images of control and treated sections were photographed under the same conditions. For figure preparation, images were exported into Photoshop CS6 (Adobe). If necessary, the levels of brightness and contrast were adjusted to the entire image. Graphics were generated using Graphpad Prism 9.0, and figures were prepared using Photoshop and InDesign CS6.

Results were processed with Graphpad Prism 9 and presented as scatter plots with mean ± SEM. Data were subjected to statistical analysis using either of the following tests, as described in the respective figure legends: Student’s t-test, Welch’s t-test, Mann-Whitney test, one-way ANOVA with post-hoc Tukey test, Kruskal-Wallis with post-hoc Dunn’s test and two-way ANOVA. All tests applied were two-tailed and a p-value ≤ 0.05 was considered significant.

## Acknowledgments

We are grateful to all the researchers whose names are mentioned along the text for generously providing reagents. We thank Dr. Eleonora Medvedev, Dr. Abed Nasereddin and Dr. Idit Shiff for technical assistance with the scRNA-seq, Tali Bdolach and Ron Rotkopf for assistance with statistical analyses and Avihu Klar for critical reading of the manuscript. G.F. is the Incumbent of the David and Stacey Cynamon Research fellow Chair in Genetics and Personalized Medicine. This study was supported by grants from the Israel Science Foundation (ISF #209/18) and from the Zelman Cowen Academic Initiatives (ZCAI) to CK.

## Author Contributions

DR performed the experiments, analyzed the data, and wrote the manuscript. GF led the bioinformatic analysis assisted by DR and CK. SO contributed to the Notch data. SK prepared probes and assisted with ISH. CK conceived and supervised the project, analyzed the data and wrote the manuscript. All authors discussed and agreed on the text and approved the manuscript.

