## Supplementary material for "Retinoic acid, an essential component of the RP organizer, promotes the spatio-temporal segregation of dorsal neural fates": Sup. Figures and legends

### Supplementary Figures and Legends

### Supplementary Figure 1

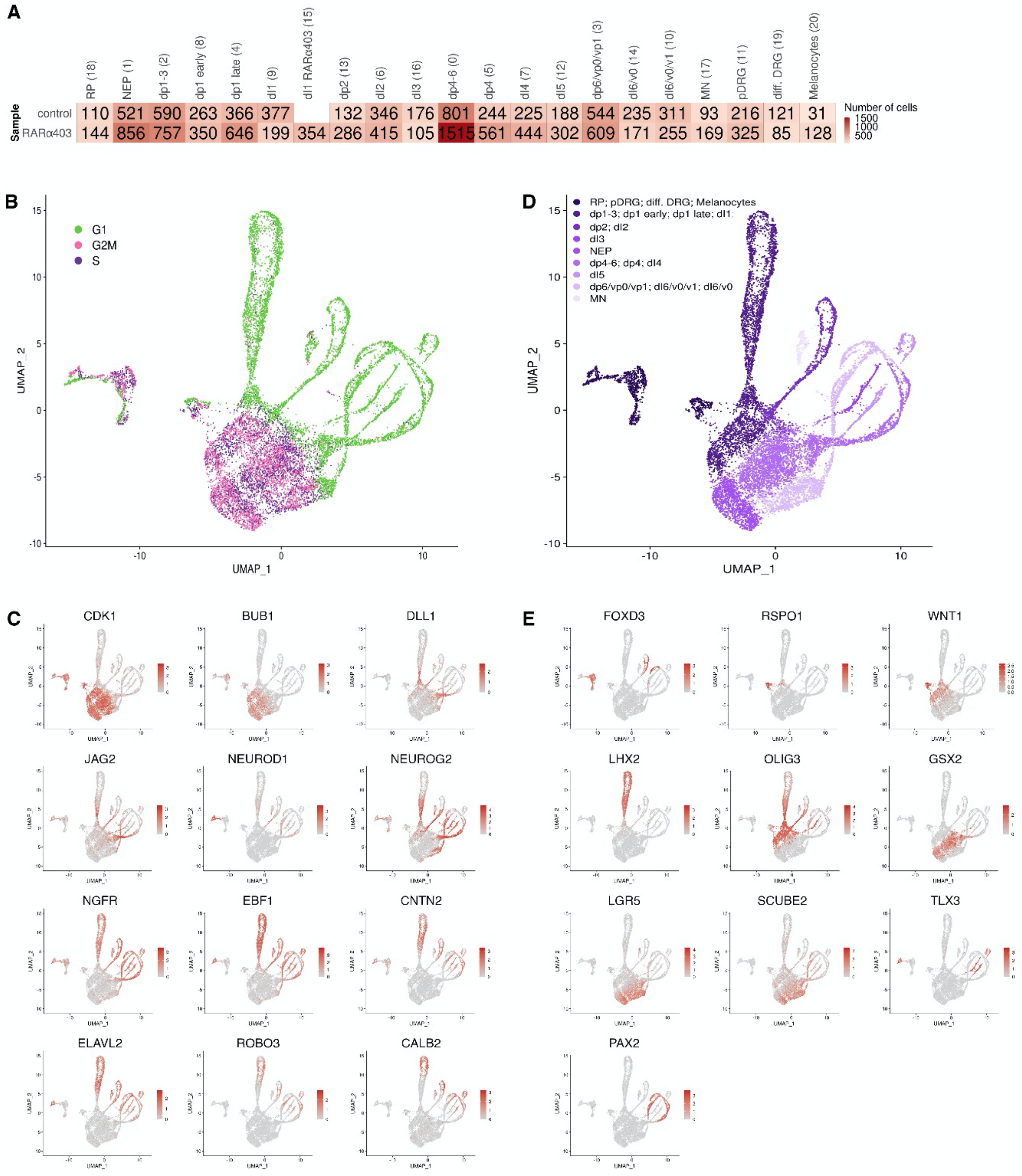


Supplementary Figure 1 – *Temporal and spatial organization of cells in the UMAP*

A) Comparison between the number of cells per cluster, in both control and RARα403 samples.

B) A UMAP colored by cell cycle phase. Note that dividing progenitors are mainly located in the octopus head. C) UMAP plots colored by the expression of selected genes corresponding to successive stages in neuronal development. D) NT cell types colored from dorsal to ventral (dark to light purple, respectively). NC derivatives are grouped with the RP. E) Selected genes that characterize various cell types in the dorsal to ventral extent of the NT.

Supplementary Figure 2

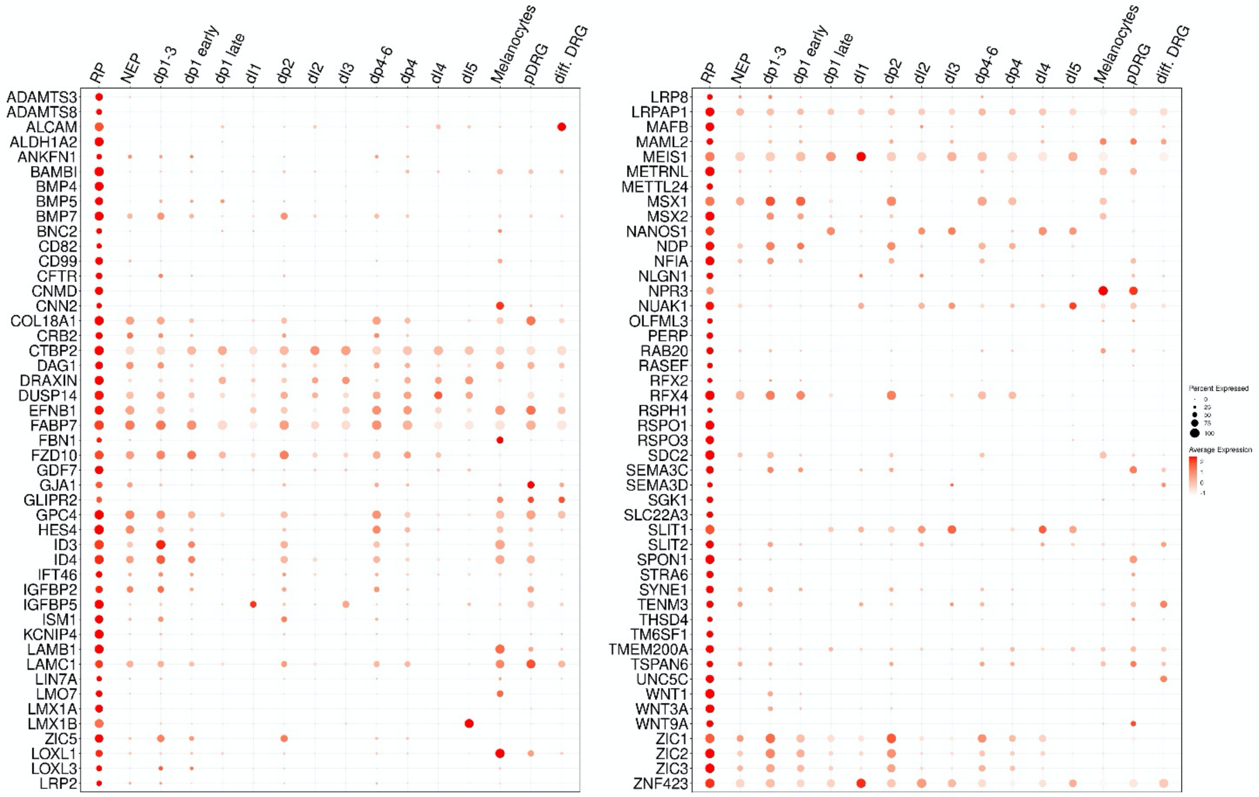


Supplementary Figure 2 – *scRNA-seq reveals an array of novel RP markers*

Dot plot of the control sample depicting expression of RP markers compared to other clusters (expression fold change >1.4 relative to other clusters).

Supplementary Figure 3

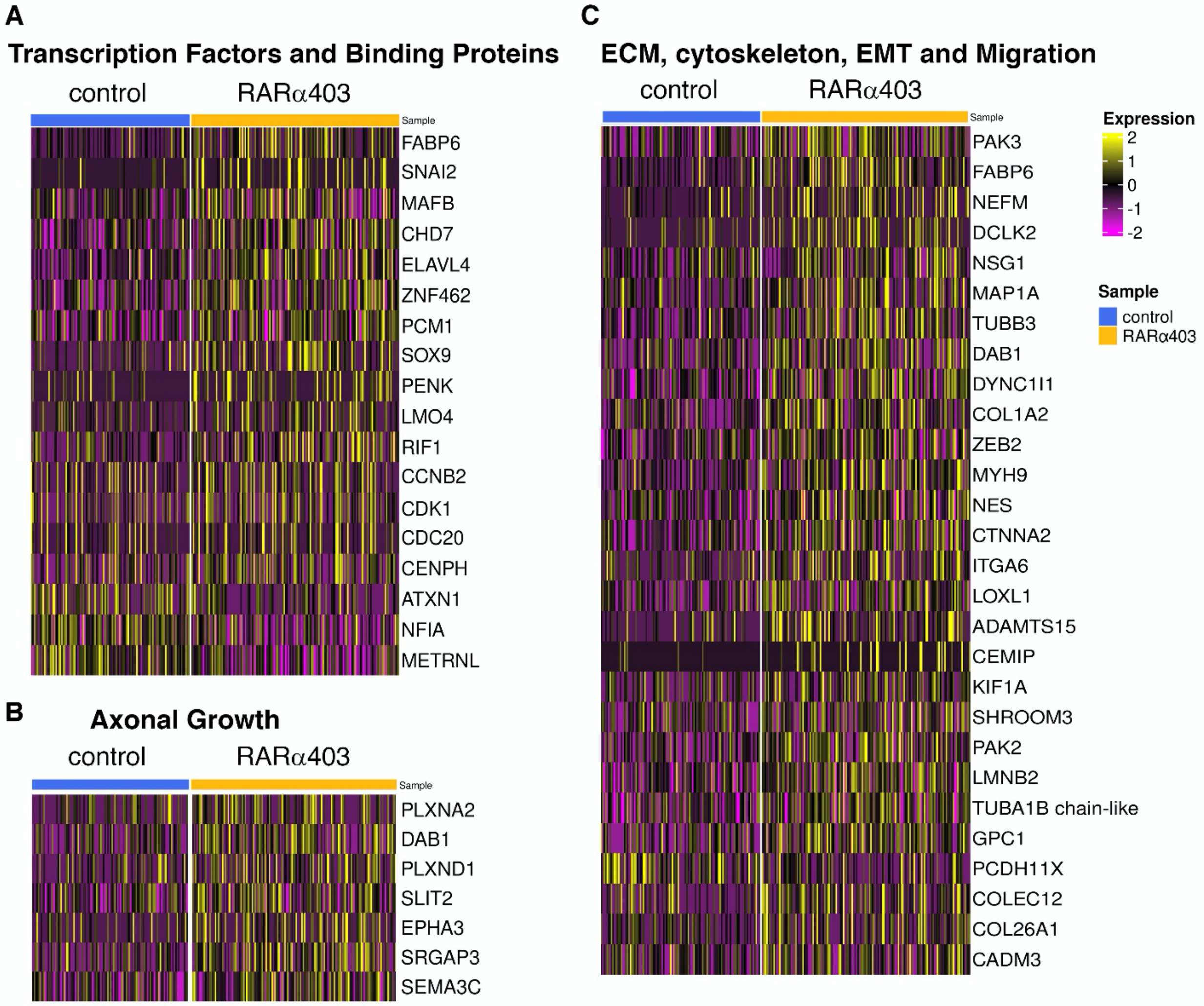


Supplementary Figure 3 - *Differential gene expression in RP reveals changes in transcription factors, adhesive properties and axonal growth*

Differential gene expression (selected from genes with minimum linear fold change ±1.3, p-value<0.05) between RARα403-treated and control RP clusters, corresponding to the following categories: A) transcription factors and binding proteins, B) Axonal growth, and C) ECM, cytoskeleton, EMT and migration.

Supplementary Figure 4

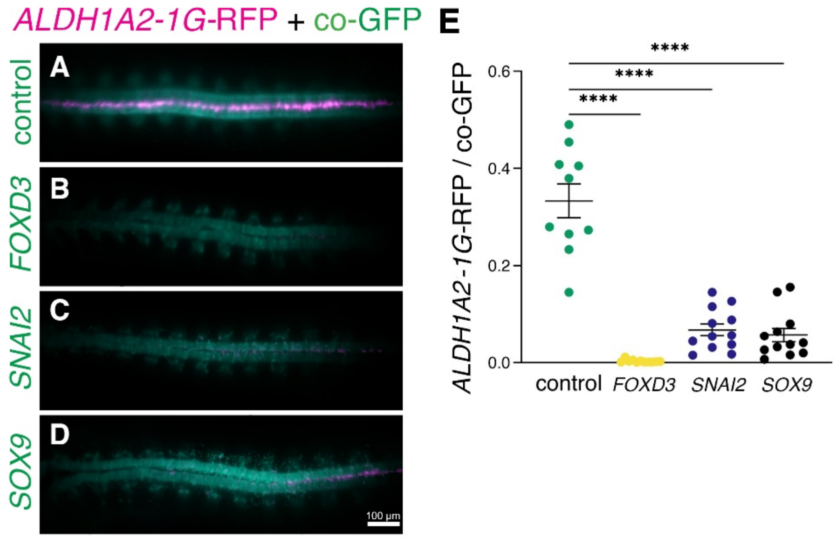


Supplementary Figure 4 – *NC and RP genes stand in a mutually repressive interaction*

Embryos were electroporated at E2.5 with either a control plasmid or a plasmid expressing *FOXD3*, *SNAI2* or *SOX9*, along with GFP to mark electroporation efficiency and location. RFP is driven by the RP-specific *ALDH1A2* enhancer. Note significant reduction of ALDH1A2 activity in treated RPs. Whole mount embryos were photographed at E4 (A-D) and intensity of RFP/GFP was quantified (E). ****$p < 0.0001$, Welch's t-test.

Supplementary Figure 5

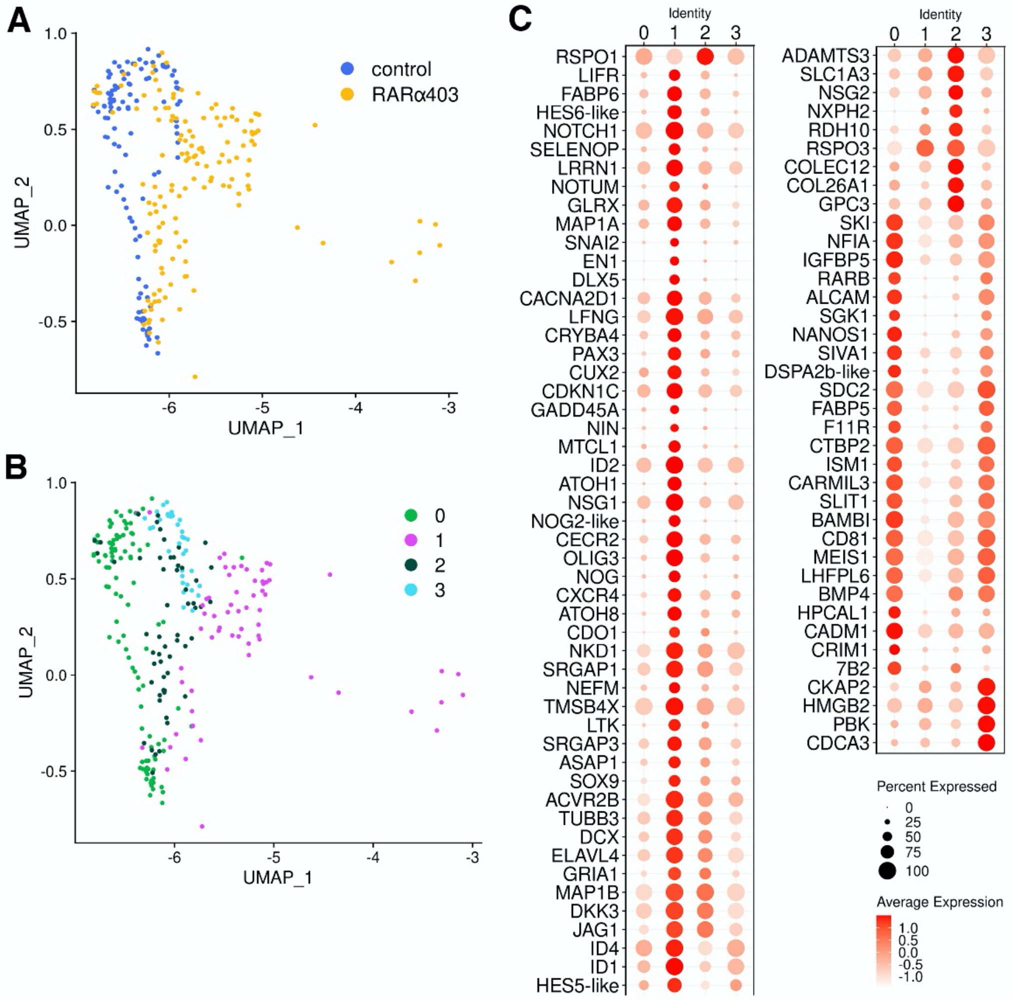


Supplementary Figure 5 – *Re-clustering of the RP reveals a subcluster with a mixed identity*

A) UMAP of the RP cluster, colored by sample. B) Re-clustering of the RP yields four subclusters. C) Dot plot depicting marker genes differentially expressed within the RP subclusters.

Supplementary Figure 6

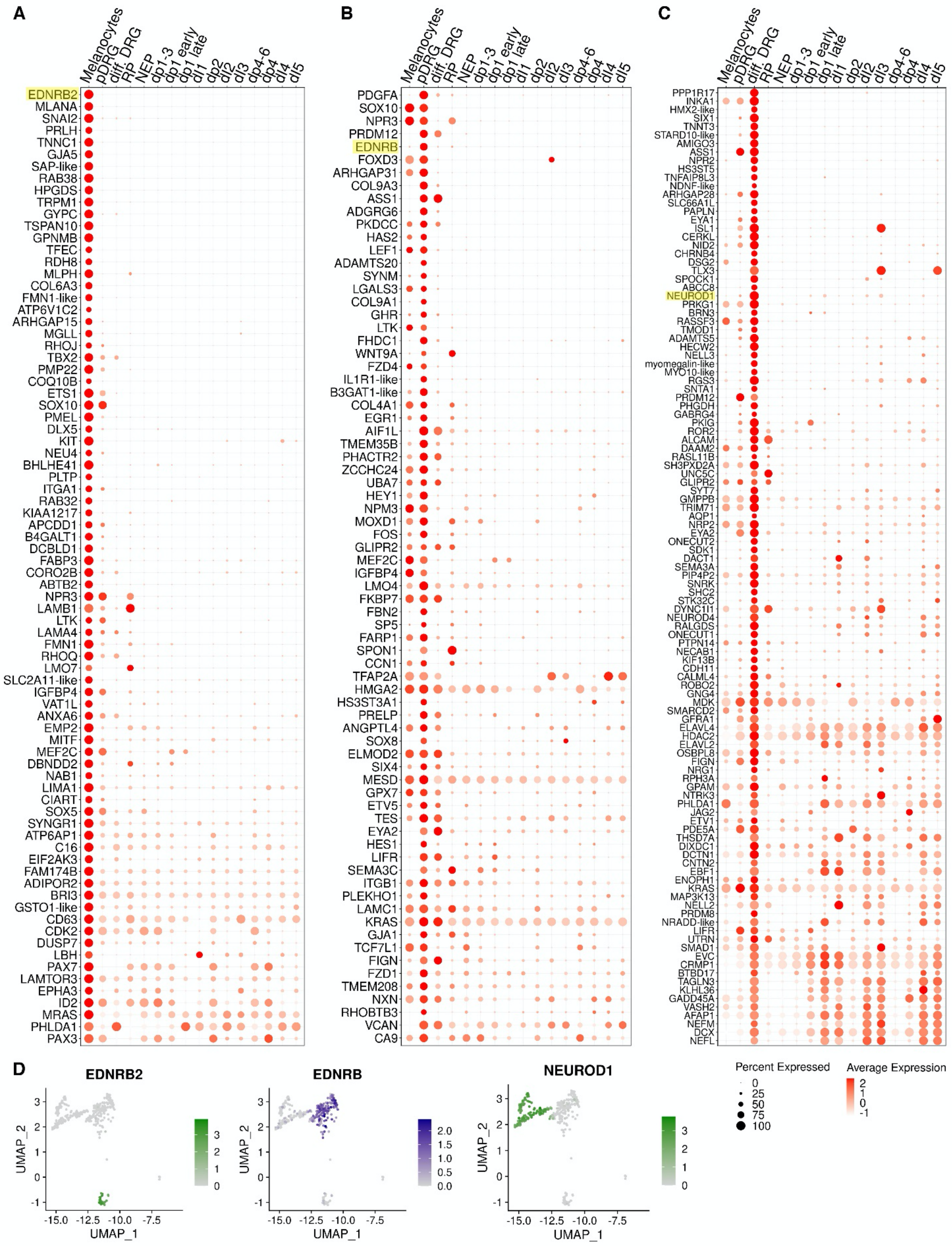

Supplementary Figure 6 – *Gene expression in NC-derived peripheral clusters*

A-C) Dot plots of the control sample depicting marker gene expression in peripheral clusters (A, melanocytes; B, pDRG; C, diff. DRG) compared to other clusters. Expression fold change >1.4 relative to other clusters. Marked in yellow are genes visualized in D. D) UMAPs demonstrating the expression of EDNRB2 and EDNRB (guidance receptors for lateral vs. ventral migration, respectively), and NEUROD1, a neuronal gene. Note the clear distinction between domains of expression.

Supplementary Figure 7

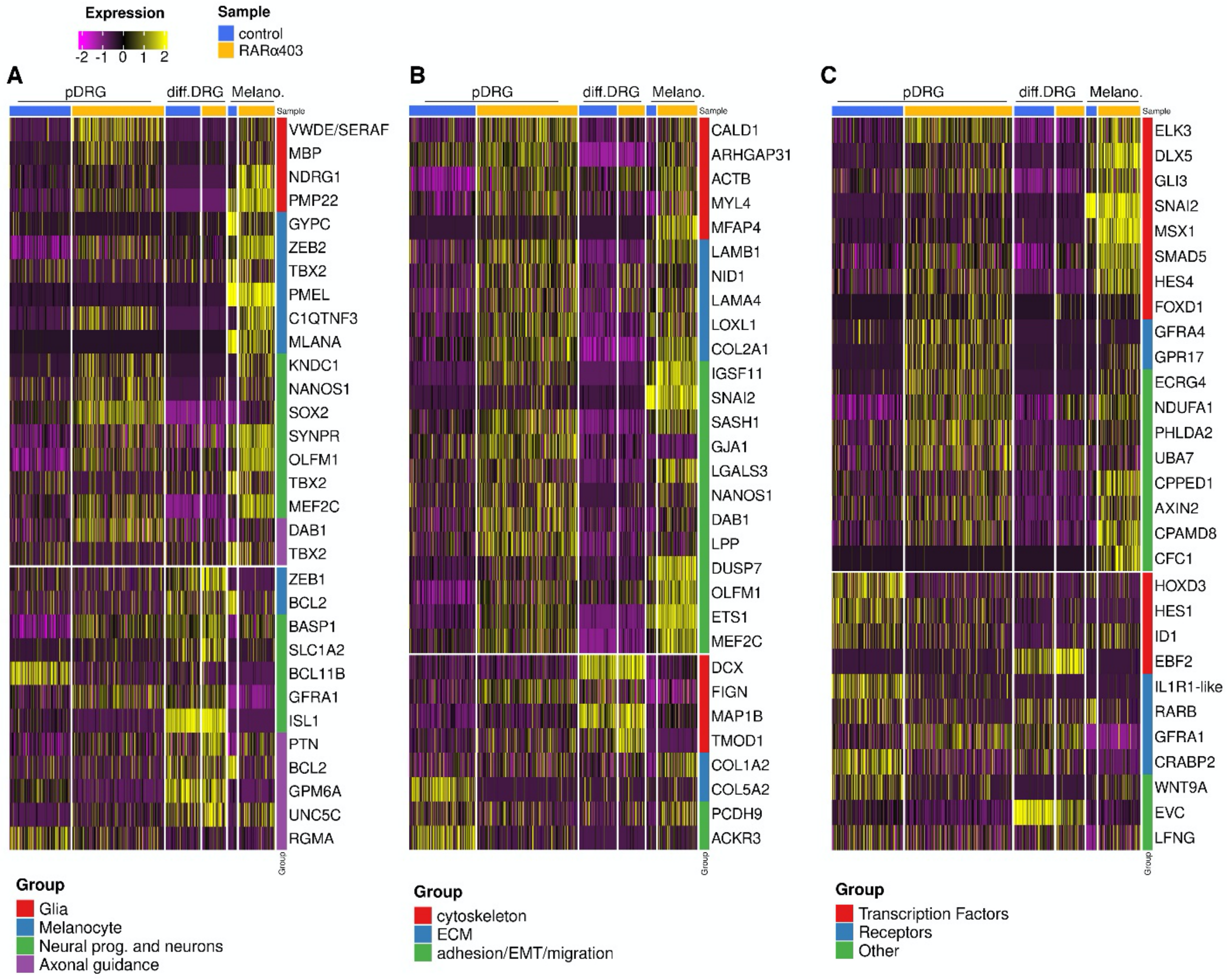


Supplementary Figure 7 – *Differential gene expression in NC-derived peripheral clusters*

Differential gene expression between RARα403-treated and control peripheral clusters, corresponding to A) Glia, melanocyte, neural and axonal guidance, B) Cytoskeleton, ECM, adhesion/EMT/migration and C) Transcription factors, receptors and other genes categories. Selected genes had a minimum linear fold change ±1.3, p-value and adjusted p-value <0.05.

Supplementary Figure 8

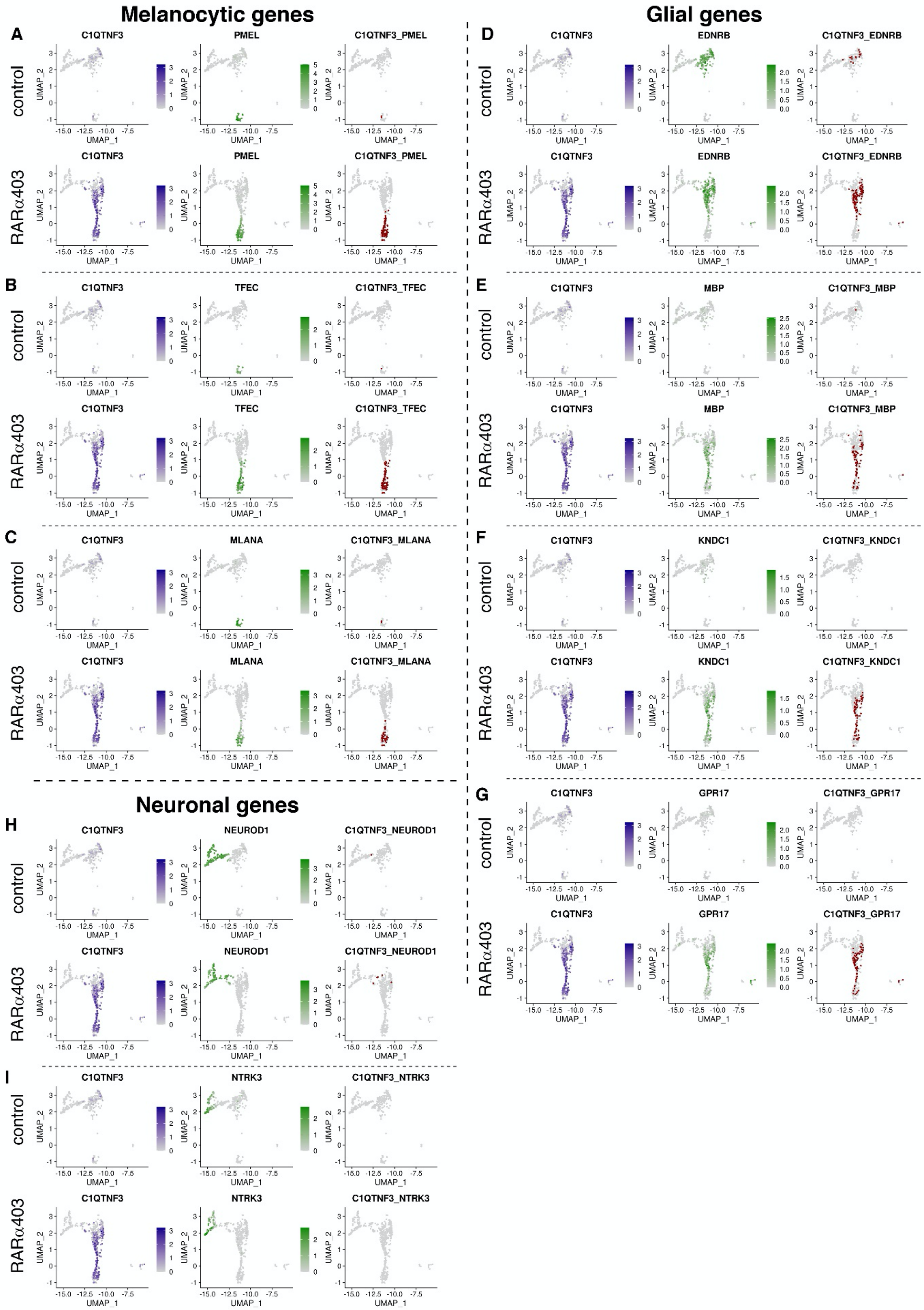

Supplementary Figure 8 – *C1QTNF3-expressing bridge cells co-express glia and melanocyte, but not neuronal genes*

Co-expression of *C1QTNF3* with melanocytic (A-C), glial (D-G) and neuronal (H-I) genes in both RARα403 and control samples, visualized on UMAPs of the peripheral clusters. Cells which had at least two reads/cell of the genes of interest are considered double-positive and stained red in the right column. A-G) Note the negligible proportion of double-positive cells in the control sample, as opposed to significant co-expression of *C1QTNF3* with glia and melanocyte genes in the bridge of the treated sample. H-I) Virtually no co-expression is detected between *C1QTNF3* and neuronal genes.

Supplementary Figure 9

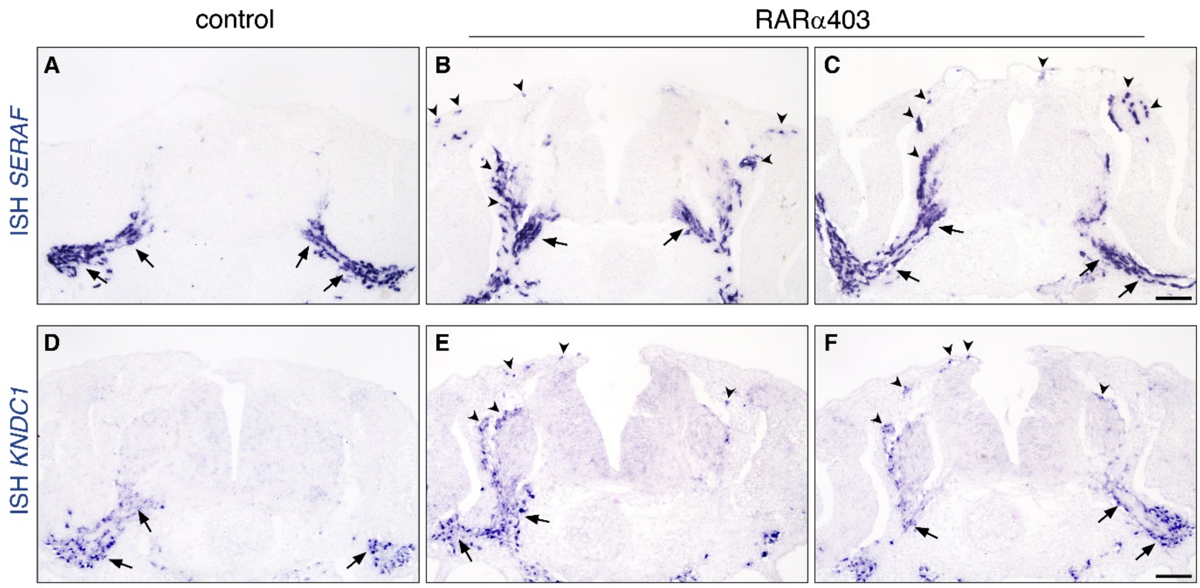


Supplementary Figure 9 – *Glial markers are ectopically expressed upon inhibition of RA activity.*

Embryos were electroporated with control PCAGG or RARα403 at E2.5, fixed at E4 and in-situ hybridized for expression of *SERAF* (A-C) or *KNDC1* (D-F). Arrows indicate normal expression sites along nerves in both control and treated embryos, whereas arrowheads highlight positive cells in the mesoderm including sites of melanocyte migration through the dermis, and DRG periphery, seen exclusively in the treated embryos (B-C, E-F). Scale bar, 50 μm.

Supplementary Figure 10

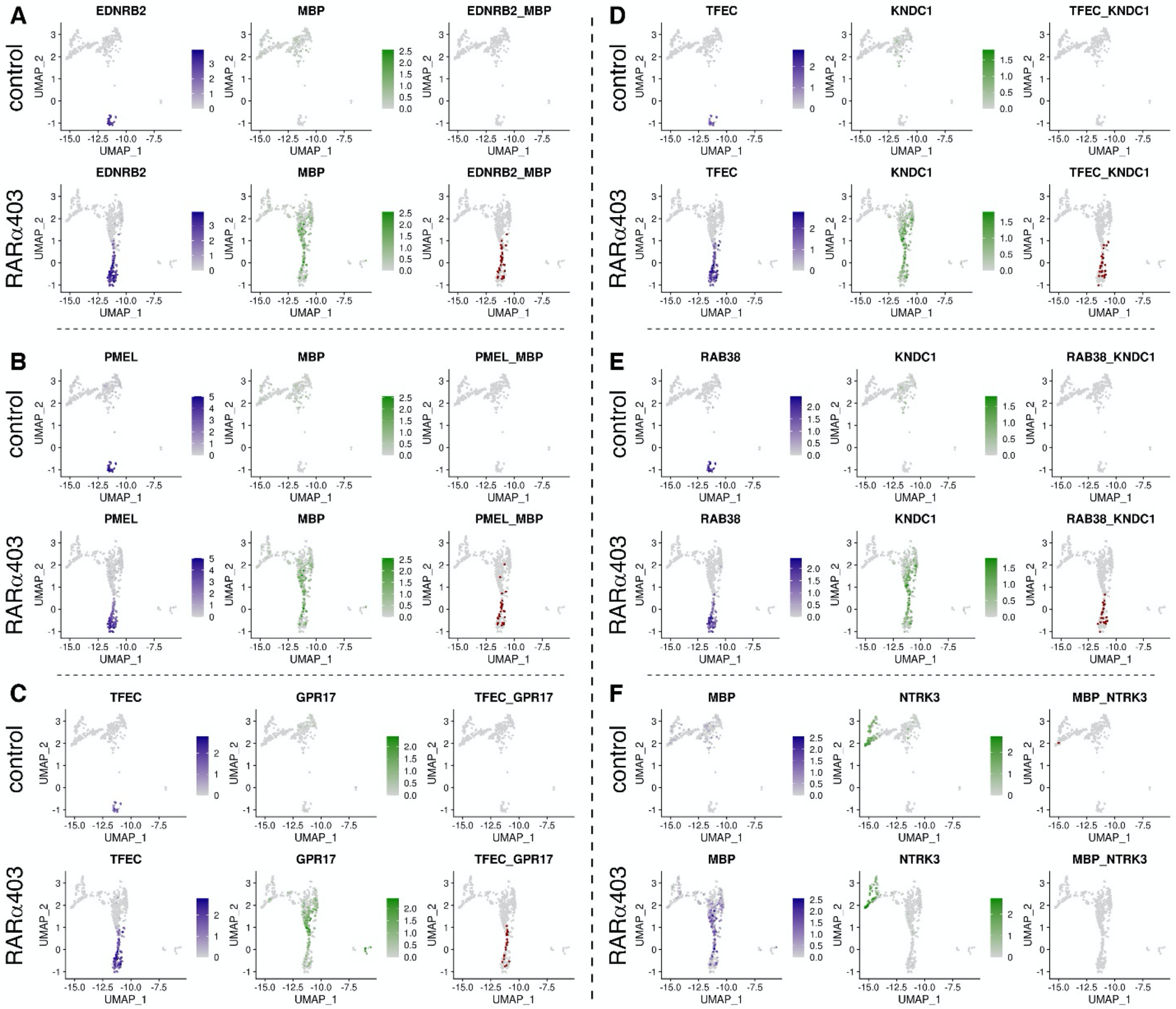


Supplementary Figure 10 – *Bridge cells co-express glia and melanocyte genes*

Co-expression of glia/melanocyte (A-E) and neuronal (F) genes in both control and RARα403 samples, visualized on UMAPs of the peripheral clusters. Cells which had at least two reads/cell of the genes of interest are considered double-positive and stained red in the right column. A-E) Note the negligible proportion of double-positive cells in the control sample, as opposed to significant co-expression of glia and melanocyte genes in the bridge of the treated sample. F) No co-expression is detected between glia (*MBP*) and neuronal (*NTRK3*) genes.

Supplementary Figure 11

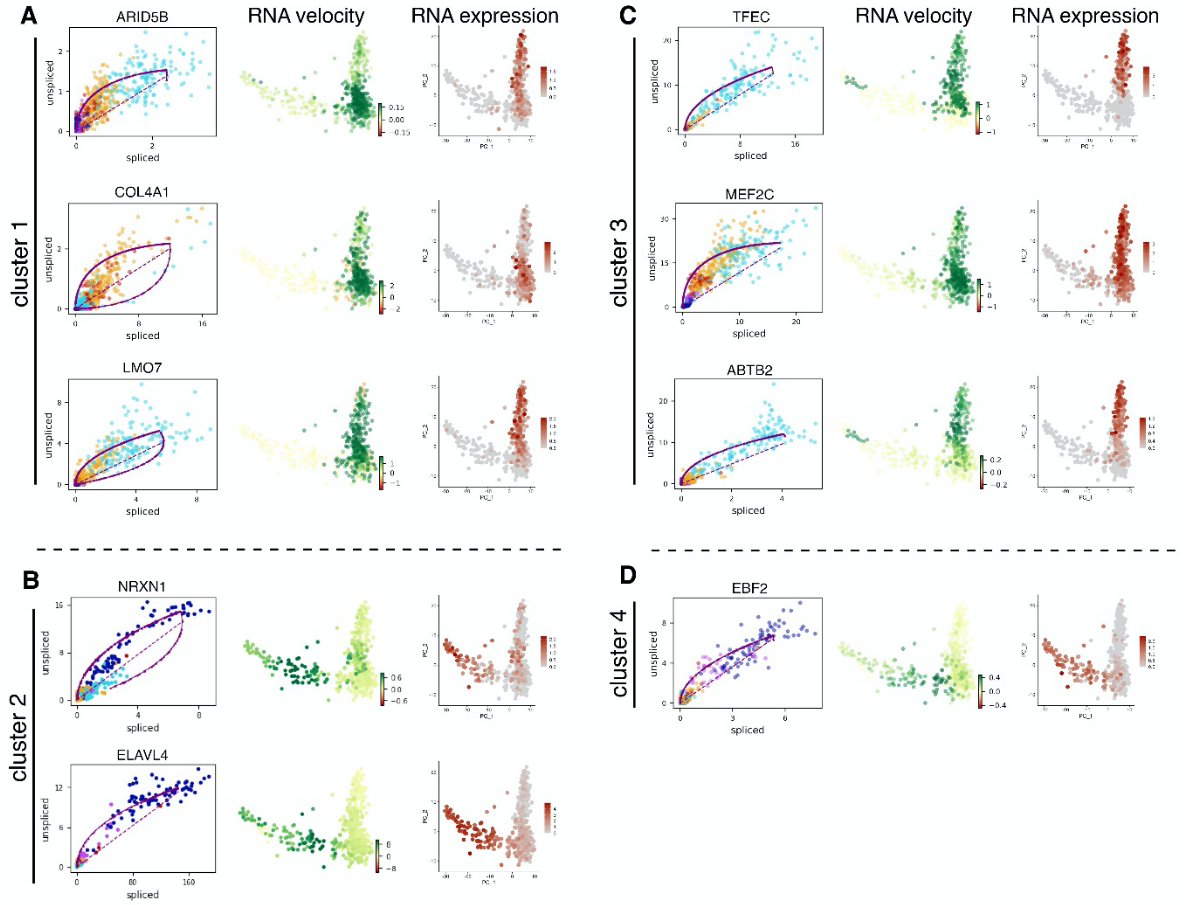


Supplementary Figure 11– *Velocity analysis reveals bridge cells as an origin of melanocytes and DRG cells*

Additional examples of specific genes with high velocity in different subclusters. Left, phase portraits of spliced versus unspliced transcripts, with a dotted line indicating the steady state of transcription and a fitted curve indicating the learned dynamics; middle, PCA colored by velocity; right, RNA expression level.

Supplementary Table 1

Table 1 – A list of identified LOC genes

| LOC | Gene |
|---|---|
| LOC107312830 | EDNRB2 |
| LOC107307993 | ZIC5 |
| LOC107317839 | HES6-like |
| LOC107308537 | SFRP2-like |
| LOC107320998 | noggin2-like/NOG2-like |
| LOC107323479 | HES5-like |
| LOC107309618 | TUBB2 chain-like |
| LOC107325741 | TUBA1B chain-like |
| LOC107310329 | FABP5 |
| LOC107314969 | 7B2 |
| LOC107314285 | DSPA2b-like |
| LOC107315152 | CECR2 |
| LOC107306834 | IL1R1-like |
| LOC107320417 | B3GAT1-like |
| LOC107313875 | HMX2-like |
| LOC107320292 | STARD10-like |
| LOC107315537 | NDNF-like |
| LOC107310125 | DSG2 |
| LOC107312758 | BRN3/POU3F3 |
| LOC107308986 | NELL3 |
| LOC107317105 | myomegalin-like |
| LOC107316896 | MYO10-like |
| LOC107312875 | GABRG4 |
| LOC107308730 | NRADD-like |
| LOC107324519 | SAP-like |
| LOC107314961 | FMN1-like |
| LOC107321228 | SLC2A11-like |
| LOC107313434 | C16 |
| LOC107309133 | VWDE/SERAF |
| LOC107316089 | KNDC1 |
| LOC107307182 | MFAP4 |
| LOC107308928 | ZEB1 |
| LOC107306284 | CFC1 |
| LOC107314568 | PHLDA2 |
| LOC107316274 | COE3-like |
| LOC107316810 | HEG1 |
| LOC107319512 | B-cadherin-like |
| LOC107324632 | PTPRV |
| LOC107317441 | SEPP1B-like |
| LOC107324454 | S100A6 |
